# Natural variation in IBF1 disrupts its interaction with CHS1 and affects metabolism of hulls in rice

**DOI:** 10.1101/2025.10.02.674788

**Authors:** Yoshiaki Ueda, Yoshinori Murata, Nozomu Sakurai, Hiroki Saito, Juan Pariasca-Tanaka, Katsuhiko Kondo, Hideki Takanashi, Takuma Ishizaki, Matthias Wissuwa

**Affiliations:** Crop, Livestock and Environment Division, Japan International Research Center for Agricultural Sciences (JIRCAS), Tsukuba, Ibaraki, Japan; Biological Resources and Post-harvest Division, JIRCAS, Tsukuba, Ibaraki, Japan; Sakura Scientific Co. Ltd., Odawara, Kanagawa, Japan; Department of Frontier Research and Development, Kazusa DNA Research Institute, Kisarazu, Chiba, Japan; Tropical Agricultural Research Front, JIRCAS, Ishigaki, Okinawa, Japan; Graduate School of Agricultural and Life Sciences, The University of Tokyo, Tokyo, Japan; Phenorob Cluster and Institute of Crop Science and Resource Conservation, University of Bonn, Bonn, Germany

## Abstract

Secondary metabolites in plants have various physiological functions, including antioxidant and antibacterial activities. Previous studies have suggested genes and associated molecular mechanisms involved in the production of diverse secondary metabolites. However, much less is known about the genetic bases underlying within-species diversity in metabolite accumulation patterns, particularly in less focused tissues such as rice hulls. In this study, we aimed to identify the causal variant that affects flavonoid accumulation in rice hulls. We identified an F-box containing protein IBF1 is causal for genotypic differences in hull color through positional cloning. The variety IR64, with straw-white hulls, harbors functional IBF1 proteins that interact with a chalcone synthase, CHS1. Conversely, frame-shift mutations of IBF1 in the variety DJ123, which has pigmented hull color, resulted in a lack of a Kelch domain essential for the IBF1-CHS1 interaction. As a result, the DJ123 variant of IBF1 (IBF1_DJ123_) no longer interacted with CHS1, which was further supported by deep learning-based protein structural modeling. Further metabolome and transcriptome analyses using IR64 and an IR64-based chromosomal segment substitution line (CSSL) carrying IBF1_DJ123_ revealed an increase in the content of multiple flavonoids (such as naringenin and luteolin), while suppressing the expression of *CAD* involved in lignin synthesis. Metabolites in the CSSL carrying IBF1_DJ123_ suppressed the growth and siderophore generation activity of *Pantoea* species, which can act as beneficial or pathogenic endophytes. This study highlights the impact of a single gene on diverse metabolite accumulation patterns and suggests that this change may provide defense against pathogens.

## Introduction

Secondary metabolites in plants have diverse physiological functions and serve as medicines and functional compounds such as insecticides and antimicrobials. Endogenous secondary metabolites also confer biotic and abiotic stress resistance to plants. For example, anthocyanins and flavonols, classes of flavonoids, protect plants from freezing stress (Schulz *et al*., 2015). Tannins induced upon mechanical damages are important flavonoids for defense against herbivores (Peters and Constabel, 2002). Accordingly, plants have developed a wide range of secondary metabolites with diverse functions, which have likely helped plants survive in natural habitats.

Secondary metabolite accumulation patterns exhibit inter- and intra-species diversity. For example, diverse tomato genotypes, including its wild relatives, showed distinct patterns of flavonoid accumulation in fruits during the ripening process (Tohge *et al*., 2020). Distinct metabolite accumulation patterns were observed among rice accessions contrasting in phosphorus use efficiency, and some secondary metabolites were suggested to serve as molecular markers for phosphorus use efficiency in rice (Watanabe *et al*., 2020). These studies highlight that the accumulation patterns of secondary metabolites are under genetic control. Several studies aimed at identifying causal genes underlying naturally-occurring genetic variations in metabolite contents. A metabolic genome-wide association study (mGWAS) using 529 rice accessions suggested sequence variations associated with genotypic differences in foliar metabolite content (Chen *et al*., 2014). Other mGWAS and large-scale metabolite profiling studies showed genetic variation in secondary metabolite content and suggested key genes for such variation in different tissues in rice (Dong *et al*., 2014; Matsuda *et al*., 2014). These studies demonstrated that the variation in metabolite content is often regulated by a small number of genetic factors, suggesting the possibility of metabolism engineering using natural sequence variation. However, such analyses were limited to certain tissues and the effect of gene modification on the metabolite profile in other tissues remains largely unknown.

Rice hulls are an important organ that physically protects seeds from mechanical and herbivorous damages. In addition to such physical functions, rice hulls contain diverse secondary metabolites that chemically protect rice seeds, with a metabolome study in rice hulls detecting > 200 flavonoids (Zhang *et al*., 2022). Momilactones are secondary metabolites derived from rice hulls that function as chemical defense against pathogens (Priego-Cubero et al., 2025). However, its content in hulls is too low to provide effective defence, and other metabolites likely play supportive roles (Yoshida *et al*., 2022). Flavonoids contained in rice hulls such as naringenin prevent the growth of pathogenic *Xanthomonas oryzae* and spore germination of *Pyricularia oryzae* (Padmavati *et al*., 1997). Similarly in wheat, flavonoids in seed coat prevent the growth of pathogenic *Fusarium* species (Skadhauge *et al*., 1997). However, knowledge on genes that increase pathogen resistance in rice hulls through metabolic engineering is still limited.

Patterns of flavonoid accumulation in rice hulls result in different types of colors, such as straw-white, purple, and brown. This can be partially explained by the *C*-*S*-*A* model in which a combination of naturally-occurring alleles at 3 loci (a bHLH transcription factor *S1*, dihydroflavonol reductase *A1*, and a MYB transcription factor *C1*) forms diverse metabolite accumulation patterns in rice hulls (Sun *et al*., 2018). Mutant and transgenic studies have also suggested genes affecting hull color in rice. A loss-of-function mutation in *CAD2*, encoding a cinnamyl alcohol dehydrogenase involved in monolignol synthesis, changes the straw-white hull color of the *indica* cultivars Z15 and H9808 to gold-brown via increased content of flavonoids (Wang et al., 2020b; Ye et al., 2024). Multiple studies have shown that mutations in INHIBITOR FOR BROWN FURROW 1 (IBF1) increase flavonoid content in hulls and change the straw-white hull of *indica* cultivars Zhefu 802, RH2B, and IR64 to brown (Shao *et al*., 2012; Xia *et al*., 2016; Li *et al*., 2025). In addition, a genome-wide association study (GWAS) using 950 diverse rice accessions also suggested that *IBF1* harbors the causal variant for hull pigmentation (Huang *et al*., 2012). These studies imply that genetic variations at *IBF1* cause changes in hull color. However, the associated molecular and structural bases and how different alleles of *IBF1* affect metabolite accumulation patterns in hulls, largely remain unknown.

During investigation, we observed that the popular *indica* variety IR64 had no pigmentation and showed straw-white hull color, while an *aus* variety DJ123 and an *indica* variety Mudgo, originating from Bangladesh and India, respectively, had strong brown pigmentation in the hull furrow (Fig. 1A). In this study, we employed positional cloning to identify the variant that causes hull pigmentation. In addition, we aimed at understanding the molecular bases related to hull pigmentation through protein-protein interaction assays and deep learning-based protein structural modeling. We further conducted metabolome analysis to evaluate the effects of causal variants on accumulation patterns of diverse metabolites. Finally, we investigated if the identified gene has a potential of increasing pathogenic resistance via altered metabolite accumulation patterns. This study reveals novel genetic and molecular bases of metabolite diversification in rice and implicates the use of hulls obtained from plants harboring different alleles in controlling microbial activities.

**Figure 1.**
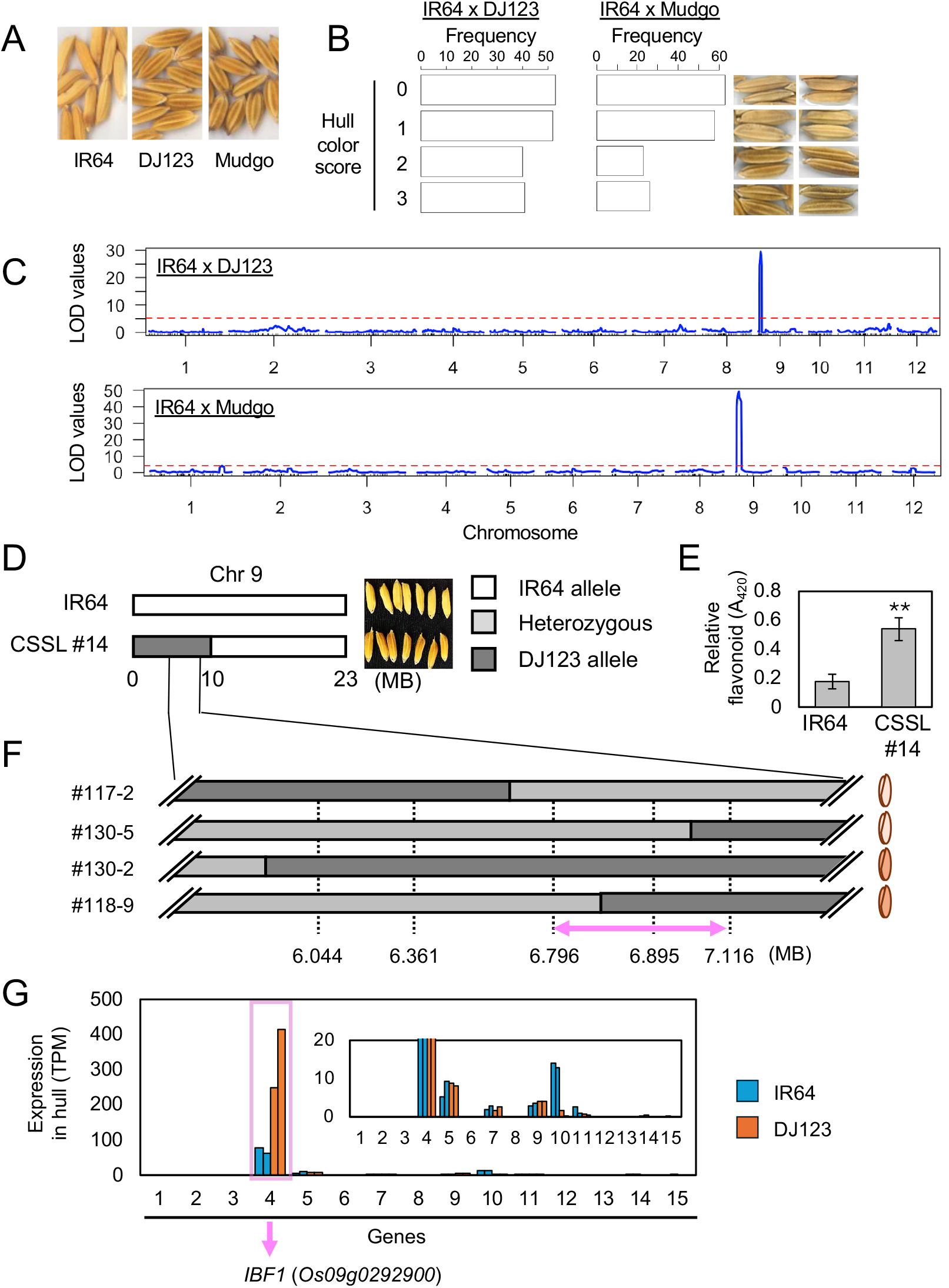
Identification of *IBF1* as the causal gene for hull pigmentation. (A) Hulls of IR64, DJ123 and Mudgo. (B) Distribution of hull color scores in two mapping populations. Hulls of representative plants with each score are shown. (C) Results of QTL mapping in two populations. The red dotted line indicates the significance threshold determined by permutation. (D,E) Genotypic description of chromosome 9 (D) and hull flavonoid content (E) in IR64 and CSSL #14. In E, values are means ± standard deviation (n=3). Student’s *t*-test was conducted, and the asterisks indicate *P* < 0.01. (F) Genotypic description of chromosome 9 QTL region in 4 fine-mapping lines and their hull color representation. The pink arrow indicates the candidate region harboring the causal gene. (G) Expression of 15 genes (transcripts per million; TPM) within the mapped region in 2 replications of IR64 and DJ123 hulls.

## Results

### Mapping of the gene responsible for hull coloration in rice

To reveal the genetic basis for hull pigmentation, we evaluated the hull color of the F_5_ population of IR64 x DJ123 recombinant inbred lines comprising 186 individuals by a score ranging from 0 (no pigmentation) to 3 (strong brown coloration) (Fig. 1B). Around half (56%) of the plants showed little or no coloration with scores of 0 or 1, while 44% had noticeable pigmentation with scores of 2 or 3 (Fig. 1B). Quantitative trait locus (QTL) mapping for this population led to the detection of a strong peak (LOD 32.0) at the 6.66 MB position on chromosome 9, flanked by markers at 6.22 MB and 7.19 MB (Fig. 1C).

We also evaluated the BC_1_F_6_ population of IR64 x Mudgo backcross inbred lines comprising 169 individuals for hull color. Around 71% of the plants had scores of 0 and 1 with little or no coloration, while 29% of plants showed strong hull coloration with scores of 2 and 3 (Fig. 1B). QTL mapping using 333 genome-wide SNP markers (Fig. S1) showed a single strong peak (LOD 49.1) at 6.70 MB on chromosome 9, flanked by markers at 2.01 MB and 7.33 MB (Fig. 1C). In both cases, the non-IR64 allele (i.e. DJ123 and Mudgo alleles) strengthened hull coloration.

We reasoned that the same genetic factor underlies the common QTL observed in both populations and carried out further characterization and fine-mapping of the QTL using the IR64 x DJ123 population. A chromosomal segment substitution line (CSSL) #14 that contained a chromosomal introgression from DJ123 at 0.0-10.1 MB region on chromosome 9 (Fig. 1D; Fig. S2) indeed exhibited a pigmented hull color, validating the result of the QTL mapping. In accordance, CSSL #14 had increased flavonoid content in hulls, which was around 3 times higher than that in IR64 (Fig. 1E). The underlying causal gene was searched by fine-mapping using BC_2_F_2_ plants of IR64 x DJ123 cross that segregate at the target locus. Hull colors of plants carrying the heterozygous and homozygous alleles at the QTL were clearly distinguished (Fig. S3), which allowed for mapping in the presence of heterozygosity. The mapping narrowed the causal region to the 6.796-7.116 MB on chromosome 9 (Fig. 1F), which contained 15 genes.

RNA-seq in young hulls of IR64 and DJ123 suggested that only 1 gene (Os09g0292900, located at 6.873 MB) out of 15 genes is highly expressed (TPM > 20) in both genotypes (Fig. 1G). This gene encodes a previously characterized F-box protein, INHIBITOR FOR BROWN HURROWS 1 (IBF1). Mutant studies showed that *IBF1* is involved in hull color pigmentation (Shao *et al*., 2012; Li *et al*., 2025). Thus, we reasoned that *IBF1* is the causal gene for hull pigmentation in DJ123 and Mudgo.

### Sequence variation at IBF1

The amino acid sequence of IBF1 in IR64 (hereafter referred to as “IBF1_IR64_”) resembled that of Nipponbare, with only 5 amino acid differences observed (Fig. S4). Prediction of protein domains suggested that IBF1_IR64_ contains an N-terminally located F-box domain and tandemly connected 3 Kelch domains (Fig. 2A) (Shao *et al*., 2012). The DJ123 type of IBF1 (hereafter referred to as “IBF1_DJ123_”) had 1-bp and 2-bp nucleotide deletions compared to IBF1_IR64_, each located directly after the F-box domain and before the 2nd Kelch domain (Fig. 2A; Fig. S4). These caused frame-shift mutations, leading to the lack of the 1st Kelch domain (Fig. 2A). The total 3- bp deletion within the coding region led to the restoration of the original reading frame towards the C-terminal, and the 2nd and 3rd Kelch domains were conserved between IBF1_IR64_ and IBF1_DJ123_ (Fig. 2A; Fig. S4).

**Figure 2.**
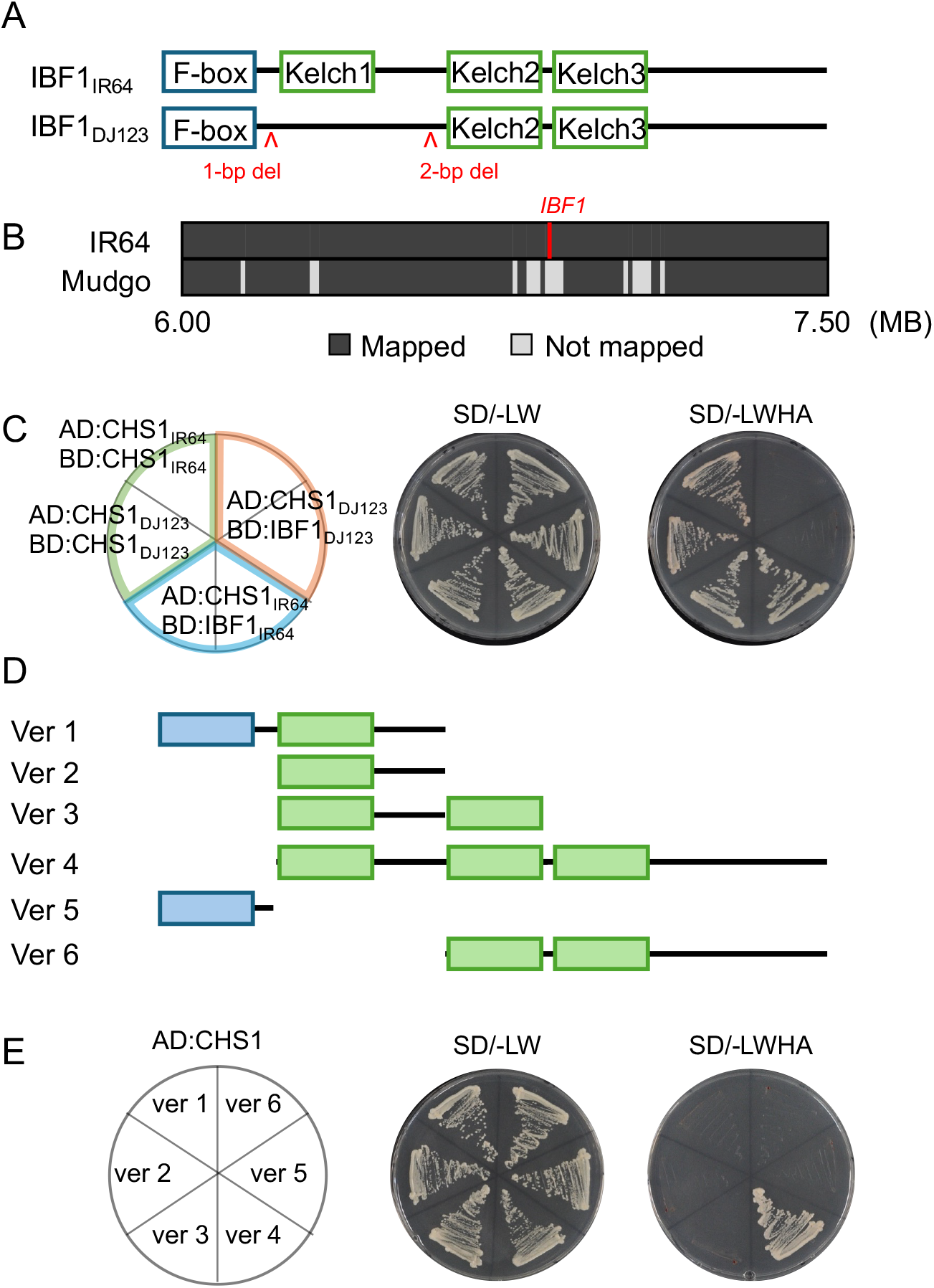
Allelic differences in IBF1 and the disruption of the IBF1-CHS1 interaction in DJ123. (A) Domain composition of IBF1_IR64_ and IBF1_DJ123_. “1-bp del” and “2-bp del” indicate nucleotide deletions in IBF1_DJ123_ compared to IBF1_IR64_. (B) Results of read alignment from whole-genome sequencing of IR64 and Mudgo to the Nipponbare reference genome near the *IBF1* locus on chromosome 9. (C) Interaction assay between IBF1 and CHS1 using a yeast two-hybrid assay. Self-interaction of CHS1 (green area) served as a positive control for the assay. (D) Truncated series of IBF1_IR64_. (E) Results of a yeast two-hybrid assay using truncated versions of IBF1_IR64_. SD/-LW and SD/-LWHA indicate selection synthetic defined (SD) media lacking Leu and Trp (-LW) or Leu, Trp, His, and Ade (-LWHA).

To analyze sequence variation near the *IBF1* locus, we carried out whole-genome re-sequencing of Mudgo. Alignment to the Nipponbare reference genome revealed that mapped reads for Mudgo were almost completely absent between the 6.869-6.907 MB region on chromosome 9, suggesting a large deletion (Fig. 2B). This result was further corroborated by PCR-based marker analyses, in which an allele-specific marker located in this region (6.895 MB) failed to amplify any fragment from Mudgo, while IR64 and DJ123 generated an amplicon for respective alleles (Fig. S5). These results suggest that a 39-kb region harboring *IBF1* is absent from Mudgo.

### IBF1_IR64_ but not IBF1_DJ123_ interacts with chalcone synthase CHS1

Plant F-box proteins interact with and affect the stability of enzymes involved in pigment synthesis (Feder *et al*., 2015; Zhang *et al*., 2017). Accordingly, a recent report in rice showed that IBF1 interacts with chalcone synthase CHS1 and promotes its degradation, thereby affecting flavonoid synthesis (Li *et al*., 2025). Thus, we hypothesized that the allele-specific mode of interaction between IBF1 and CHS1 explains the phenotypic difference in hull color and conducted yeast two-hybrid (Y2H) assays. In this assay, the self-interaction of CHS1 (Waki *et al*., 2020) served as a positive control (Fig. 2C, green area). We confirmed the absence of yeast growth on the selection media (SD/-LWHA) in the absence of the interacting partner (Fig. S6). The assay showed that IBF1_IR64_ interacted with CHS1 (Fig. 2C, blue area), consistent with the observation in a genotype showing straw-white hull color (Li *et al*., 2025). Y2H assays using a series of truncated IBF1_IR64_ suggested that the F-box domain was not necessary for the IBF1-CHS interaction (Fig. 2D,E), supporting the results of a previous study (Li *et al*., 2025). On the other hand, the lack of either the 1st or 3rd Kelch domain in IBF1_IR64_ impaired the IBF1-CHS1 interaction (Fig. 2D,E). Consistent with the lack of the 1st Kelch domain, IBF1_DJ123_ did not interact with CHS1 (Fig. 2C, orange area).

We investigated potential interactions of IBF1 with phenylalanine ammonia lyases (PALs), another class of putative targets of F-box proteins (Zhang et al., 2013; Zhang et al., 2015). Apart from *CHS1, PAL2/4/5/7* were abundantly expressed (TPM > 50) in hulls of both IR64 and DJ123 (Fig. S7A), and the analysis was limited to these 4 PALs. The Y2H assay showed that neither IBF1_IR64_ nor IBF1_DJ123_ interacted with PAL2/4/5/7 (Fig. S7B), corroborating that the mode of interaction between IBF1 and PAL is not causal for the difference in hull pigmentation.

### Structural basis for the IBF1-CHS1 interaction

To understand the structural basis of the IBF1_IR64_-CHS1 interaction, the 3D structure of the complex was modeled using the deep learning-based protein complex structure prediction tool Alphafold3 (Abramson *et al*., 2024). The structure of the IBF1_IR64_-CHS1_IR64_ complex was confidently modeled, with overall pIDDT values mostly exceeding 70, indicating high accuracy of local structure (Fig. 3A). Additionally, high ipTM (0.74) and pTM (0.81) values ensured a confident prediction of the overall structure and domain-to-domain relationships. The three Kelch domains of IBF1 formed a ring-shaped structure called the β-propeller domain (Gray *et al*., 2009), potentially serving as the interaction surface with CHS1 (Fig. 3B). Consistent with the Y2H assays, CHS1 had high predicted aligned error (PAE) with the F-box domain of IBF1_IR64_ but low PAE with the 1st Kelch domain onwards, suggesting the importance of the Kelch domains for the interaction with CHS1 (Fig. 3C). Prediction of critical residues for the interaction suggested that 86 pairs of residues between IBF1_IR64_ and CHS1_IR64_ may mediate the interaction. In contrast, the accuracy of the IBF1_DJ123_-CHS1_DJ123_ complex was low, with a large region having low (< 50) pIDDT and relatively low ipTM (0.62) and pTM (0.71) values (Fig. 3D). The lack of the 1st Kelch domain in IBF1_DJ123_ disrupted the interaction surface, forming an incomplete propeller shape (Fig. 3E). The PAE was also high for the region before the 2nd Kelch domain (Fig. 4E). Prediction of critical residues indicated that only 55 pairs of residues may mediate the IBF1_DJ123_-CHS1_DJ123_ interaction, consistent with the observation that they do not form a stable complex. Similar results were obtained using IBF1_DJ123_ and CHS1_IR64_ (Fig. S8).

**Figure 3.**
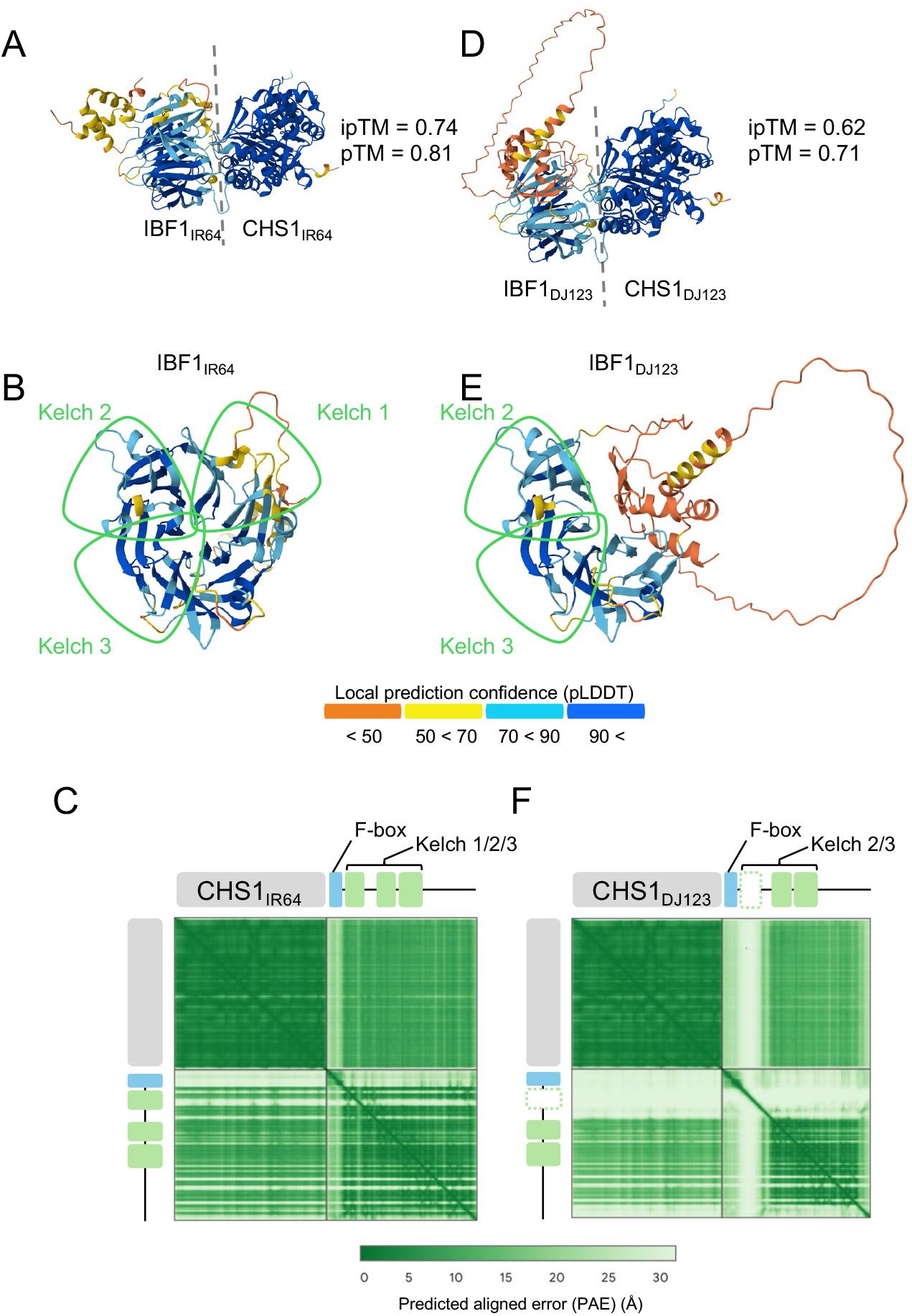
3D structure modeling of the IBF1-CHS1 interaction. (A,D) Structural modeling for IBF1_IR64_-CHS1_IR64_ (A) and IBF1_DJ123_-CHS1_DJ123_ (D). (B,E) The interaction surface of IBF1_IR64_ (B) and IBF1_DJ123_ (E) viewed from the CHS1 side. The CHS1 molecule is omitted from visualization for simplicity. In A, B, D and E, amino acid residues are colored by the accuracy level of prediction judged by pIDDT values. (C,F) Expected position error for the modeling results for IBF1_IR64_-CHS1_IR64_ (C) and IBF1_DJ123_-CHS1_DJ123_ (F).

**Figure 4.**
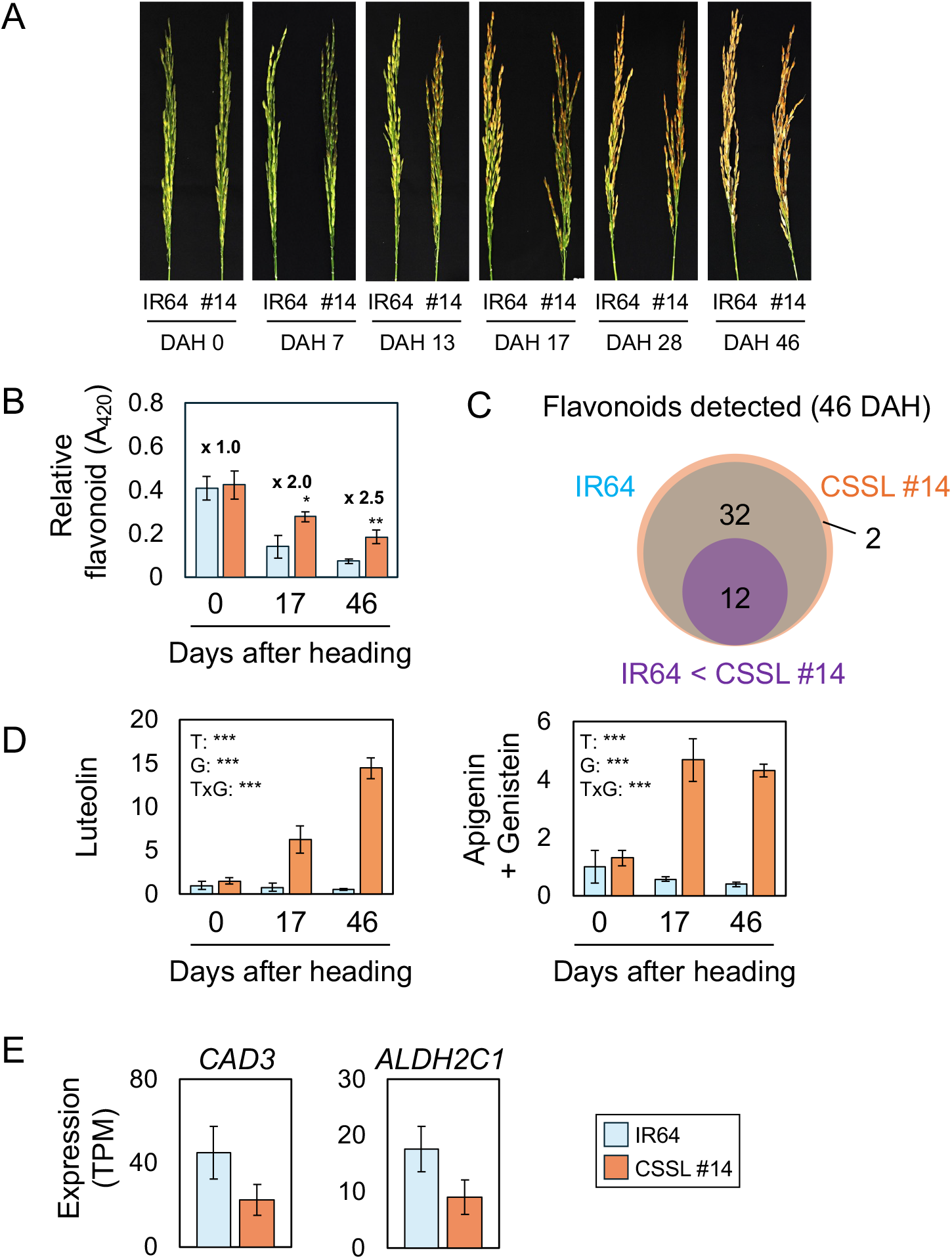
Effects of IBF1 variants on metabolite accumulation and gene expression patterns at different time points. (A) Appearance of panicles of IR64 and CSSL #14 at different time points in the field. DAH, days after heading. (B) Time-course analysis of flavonoid content in hulls of IR64 and CSSL #14. (C) Results of untargeted metabolome analysis using hull samples from 46 DAH. (D) Relative abundance of selected metabolites in hull samples from 0, 17, and 46 DAH. (E) Results of RNA-Seq using hull samples from 17 DAH. In B, D and E, values are means ± S.D. (n=3 for B, D and E). In B, Student’s *t*-test was conducted for each time point, and the results are shown as asterisks (**, *P* < 0.01; ***, *P* < 0.001). In D, Two-way ANOVA was conducted for each metabolite to analyze the effect of time (T), genotype (G), and their interaction (TxG), and the results are shown as asterisks (***) for *P* < 0.001.

Previous studies have identified IBF1 mutations that cause hull pigmentation, including G338D (Xia *et al*., 2016) and V340D (Li *et al*., 2025) mutations, with another mutation, R374C, being suggested without experimental validation (Li *et al*., 2025). Protein structural analysis suggested that the G338D mutation disrupted a part of the loop structures (Fig. S9A), which eventually disrupted the interaction surface originally formed in the wildtype version of IBF1_IR64_ (Fig. S9B). The R374C mutation was also predicted to cause a disruption of the interaction surface (Fig. S9C). The effect of the V340D mutation was less clear, but it partially affected the loop structure near the interaction surface (Fig. S9D

### Effects of allelic difference of IBF1 on metabolic profile

The analysis of hulls at different developmental stages indicated that the effect of IBF1 was near-absent immediately after heading, but differences in hull color became evident by 7 days after heading (DAH), which further increased towards maturation (Fig. 4A). Total flavonoid content in hulls was quantified at different time points using the Al^3+^-chelation method (Mammen and Daniel, 2012). We note that the 80% methanol extracts from 0 and 17 DAH were colored even in the absence of Al^3+^ (Fig. S10), and mixture with Al^3+^ solution led to the formation of precipitates, as reported for plant samples containing chlorophyll (Chen et al., 2001). Thus, the flavonoid content for 0 and 17 DAH may have been overestimated, but the comparison between IR64 and CSSL #14 followed a trend similar to the visual observation of the panicles: the flavonoid content in CSSL #14 was comparable with IR64 immediately after heading (0 DAH), but it drastically increased by 17 DAH. By 46 DAH, flavonoids in CSSL #14 reached levels 2.5 times higher compared to IR64 (Fig. 4B).

To analyze global metabolic changes caused by different alleles of IBF1, we conducted an untargeted metabolome analysis using hulls of IR64 and CSSL #14 at complete maturation (46 DAH). Among a total of 7,994 detected metabolite peaks in positive mode, 1,625 and 1,806 were only detected or more abundant (i.e. > 3-fold difference) in IR64 and CSSL #14, respectively (Table S1). Prediction of unannotated metabolites by FlavonoidSearch (Akimoto *et al*., 2017) using mass-spectrum data for 744 metabolite peaks suggested the detection of a total of 46 flavonoids, including one that was likely automatically or artificially generated by fragmentation of its derivative (glycoside) (see Supplemental Note 1). Among the candidate 46 flavonoids, 2 and 14 were detected only in CSSL #14 and absent in IR64. None of the predicted flavonoids was more abundant in IR64 (Fig. 4C; Table S2). Quantification results of confirmed metabolite peaks showed that flavonoids such as apigenin/genistein (undistinguishable) and luteolin were more abundant in CSSL #14. Prediction of unannotated peaks using MASST (Wang et al., 2020a) and MS-FINDER (Tsugawa et al., 2016) software suggested that one of the metabolites unique in CSSL #14 was tricin. Similarly, one of the metabolites more abundant in CSSL #14 (6.3 times more abundant) was predicted to be naringenin (Table S2).

Selected metabolites were quantified in a targeted manner with replications using hull samples harvested at 0, 17 and 46 DAH. We note that the heading date of IR64 and CSSL #14 was the same (71 days after transplanting), excluding the bias caused by the difference in the developmental stage. The content of luteolin was almost constant in IR64 during the hull development, while CSSL #14 accumulated a much higher amount, particularly at the later time points (Fig. 4D). Apigenin/genistein also followed a similar trend (Fig. 4D). The content of Tyr, Phe, Ile, and Leu was not significantly different between IR64 and CSSL #14, with the content markedly decreasing over time in both genotypes (Fig. S11).

### Effects of allelic differences of IBF1 on gene expression profile in hulls

RNA-Seq revealed that 78 and 42 genes were upregulated in IR64 and CSSL #14 hulls at 17 DAH, when the flavonoid content in CSSL #14 started to increase compared to IR64 (Table S3). Genes upregulated in IR64 included *CINNAMYL ALCOHOL DEHYDROGENASE 3* (*CAD3*) (Os10g0430200) involved in lignin synthesis, and *ALDEHYDE DEHYDROGENASE 2C1* (*ALDH2C1*) (Os01g0591300) involved in the generation of lignin precursors (ferulate and sinapate) (Fig. 4E). Additionally, a search on the KEGG database (https://www.genome.jp/kegg/kegg2.html) (Kanehisa et al., 2025) revealed that no genes encoding flavonoid synthesis enzymes were differentially expressed between IR64 and CSSL #14 hulls (Table S3).

### Effects of allelic variation of IBF1 on microbial activities

The above analyses indicated that different alleles of *IBF1* drastically change the metabolic profile of rice hulls. Since secondary metabolites play a crucial role in the physiological functions of hulls, we examined the impact of different *IBF1* alleles on microbial activity in hull extracts. We selected two strains of *Pantoea* species (*Pantoea agglomerans* strain 102470 and *Pantoea* species strain 6PNS2) as model microbe species due to their agricultural significance as endophytic plant pathogens and beneficial microbes (Dutkiewicz et al., 2016). The hull extracts from CSSL #14 used in the experiment had a higher total flavonoid content (A_420_ = 0.047 and 0.074 for IR64 and CSSL #14, respectively). The addition of hull extracts from IR64 and CSSL #14 did not significantly affect the cellulase and phosphorus solubilization activities of either strain (Fig. 5A). However, the siderophore producing activities, which enhance iron acquisition through chelation, were notably decreased when hull extracts from CSSL #14, but not from IR64, were added to the growth medium in both strains (Fig. 5A,B). The hull extracts from CSSL #14 inhibited the growth of strain 102470, while those from IR64 did not (Fig. 5C). Similar analyses using *Staphylococcus* species (*S. pasteuri* strain 105 and *S. warneri* strain 108) showed no significant effect of hull extracts on physiological parameters (cellulase, phosphorus hydrolysis and siderophore production activities). However, hull extracts of CSSL #14 did suppress growth of *S. warneri* strain 108 (Fig. S12).

**Figure 5.**
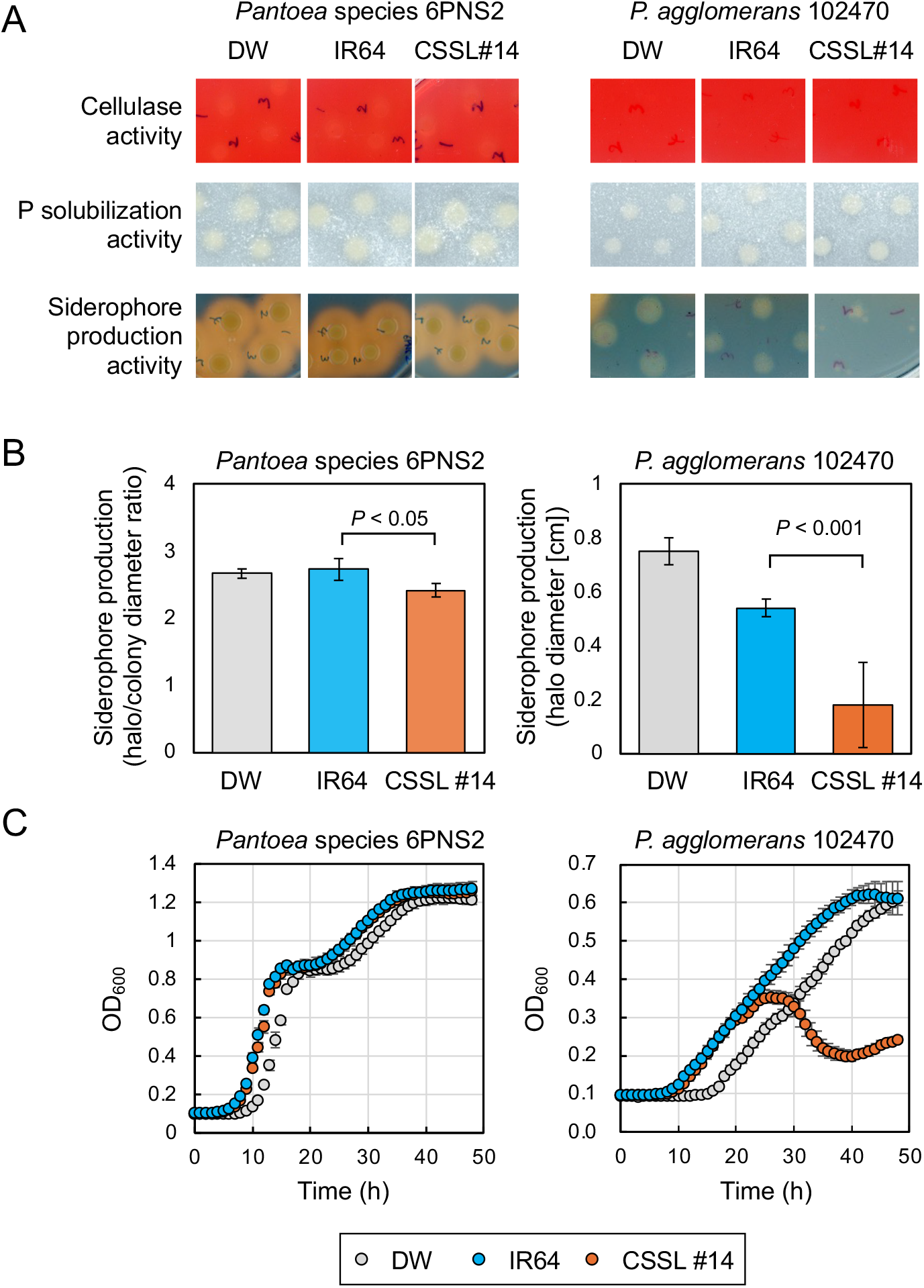
Effects of hull extracts from IR64 and CSSL #14 on physiological activities and growth of microbes. (A) Analysis of physiological activities through plate-based assays using different substrates. (B) Quantitative analysis of siderophore producing activity under the conditions in A. Note that the ratio of halo to colony diameter was quantified for *Pantoea* strain 6PNS2, while the halo diameter was quantified for *P. agglomerans* 102470 since colony growth was not observed. (C) Growth analysis. In B and C, values are means ± S.D. (n=3 and 6 for DW and hull extracts in B, n=6-8 in C). In B, Student’s *t*-test was conducted to compare the values obtained using the hull extracts of IR64 and CSSL #14, and the resultant *P*-values are shown.

### Distribution of alleles critical for pigmentation in wider genetic resources

The above analysis suggested that the 1-bp and 2-bp nucleotide deletions in *IBF1* are crucial for the loss of the first Kelch domain and the loss of IBF1-CHS1 interaction, ultimately leading to hull pigmentation. We examined the distribution of these critical nucleotide deletions in a broader population of rice. Looking at the presence of these nucleotide deletions among 3,000 rice accessions (Wang et al., 2018), we found that the 1-bp and 2-bp deletions were only detected in 14 and 10 accessions, resulting in the allele frequencies of 0.004 and 0.003, respectively. Most of these deletions were identified within the *aus* subpopulation, with 57% and 80% of the lines carrying the 1-bp and 2-bp deletions falling into the *aus* subpopulation, while the remaining lines were classified as the *indica* subpopulation (Table S4). All the lines with these mutations originated from South Asia (i.e. Bangladesh and India), except for one line with an unknown origin (Table S4). Among the 20 accessions carrying at least one of the 1-bp and 2-bp deletions, all 12 lines with available hull color data had pigmented hulls, including both brown and purple-black hulls (Table S4).

## Discussion

Flavonoids play multifaceted roles, such as protection against herbivores and tolerance to abiotic stress including high light and cold stress. Genotypic differences in flavonoid accumulation patterns likely reflect the diverse environments in which plants have evolved. Despite recent elucidation of metabolic profiles and physiological functions of each metabolite, much less is known about the genetic bases for genotypic differences in metabolite accumulation and functionality of plant extracts, particularly in less focused tissues like rice hulls. In this study, we focused on the differences in hull color in rice and identified the causal variants, associated molecular bases, and functional consequences.

### Changes in metabolic patterns caused by the interaction between F-box proteins and metabolic enzymes

We have shown that either frame-shift mutations (DJ123) or the absence (Mudgo) of *IBF1* are the causal variants that intensify hull pigmentation. In genotypes with straw-white hull color, functional IBF1 actively degrades CHS1 (Li *et al*., 2025), while absence or dysfunction of IBF1 likely maintains CHS1 more intact in DJ123 and Mudgo, accelerating flavonoid synthesis. This molecular mechanism is consistent with an Arabidopsis Kelch repeat F-box protein, KBF^CHS^, which specifically interacts and destabilizes CHS, affecting anthocyanin accumulation and tissue pigmentation (Zhang *et al*., 2017). A similar mechanism was proposed in muskmelon, with a Kelch domain-containing F-box protein, CmKFB, suggested as the key factor for genotypic differences in flavonoid accumulation and fruit rind color (Feder *et al*., 2015). On the other hand, homologues of Arabidopsis KBF^CHS^, KFB01/20/39/50, exert their influence by interacting with and destabilizing PALs (Zhang et al., 2013; Zhang et al., 2015). In rice, a Kelch-repeat F-box protein, OsFBK1, interacts with cinnamoyl CoA reductase (CCR) that catalyzes the first step of lignin synthesis and affects anther and root cell wall thickening (Borah and Khurana, 2018). The same study also showed that the lack of a Kelch domain in OsFBK1 impairs its interaction with CCR, supporting our model that a lack of one Kelch domain was sufficient to affect the phenotype. Thus, different groups of Kelch domain-containing F-box proteins have different interacting partners to exert their influence, probably by forming various interaction surfaces formed by multiple Kelch domains (Borah and Khurana, 2018). Considering this, it is plausible that rice homologues of IBF1, likely those expressed in other tissues, interact with PALs and affect tissue pigmentation through altered stability of PALs. Further studies are necessary to comprehensively elucidate the interacting partners for other Kelch domain-containing F-box proteins, as well as identifying the key amino acid residues determining the specificity of the interaction.

### Changes in metabolic patterns in rice hulls caused by different alleles of IBF1

CHS generates naringenin chalcone from *p*-coumaroyl CoA, which is otherwise used to biosynthesize aldehydes that serve as precursors for lignin synthesis (Lam *et al*., 2021). Thus, CHS1 functions as a crucial gate in controlling the metabolic flow. This is further supported by the observations that differentially expressed genes in IR64 and CSSL #14 did not contain any known genes encoding flavonoid synthesis enzymes (Table S3) and that no flavonoids were more abundant in IR64 (Table S2). These strongly suggest that the difference in flavonoid metabolism between IR64 and CSSL #14 is not caused by the difference in the expression of flavonoid synthesis genes but by the supply levels of the substrate. This mode of action in IBF1 is in contrast to *C1*, which changes metabolism primarily via altered expression of flavonoid metabolism genes (Sun et al., 2018).

The mechanism proposed above also explains differences in the transcriptome between IR64 and CSSL #14. The substrate of CHS, *p*-coumaroyl CoA, is also used to synthesize *p*-coumaraldehyde, sinapaldehyde and coniferaldehyde, all of which are used for lignin synthesis (Lam *et al*., 2021). CAD converts these aldehydes to generate *p*-coumaroyl alcohol, sinapyl alcohol, and coniferyl alcohol, which are used as building blocks for H-, S- and G-lignins, respectively. The higher expression of *CAD3* in IR64 (Table S3) may indicate that aldehydes derived from *p*-coumaroyl CoA are more abundant in IR64 since they are not used for flavonoid synthesis, and that lignin synthesis is more active in IR64.

These findings suggest that the synthesis of flavonoids and lignin could be in a trade-off relationship. A rice mutant carrying an amino acid substitution in *CAD2* had 20% lower lignin and 40-50% higher flavonoid content in hulls, leading to hull pigmentation (Wang et al., 2020b). On the other hand, foliar lignin content was not increased in a rice mutant lacking CHS1, despite a significant decrease in contents of flavonoids such as apigenin and luteolin (Lam *et al*., 2021). Thus, the relationship between lignin and flavonoid synthesis is not straightforward, probably because some flavonoids such as tricin, naringenin and apigenin are incorporated into lignin in rice (Lam *et al*., 2021). Further investigations are necessary to determine if different alleles of IBF1 affect the content and composition of lignin and thus affect the mechanical properties of hulls, which may be linked to resilience to herbivore damage.

### Structural basis of dysfunctional IBF1 that cause hull pigmentation

Our 3D modeling using Alphafold3 suggested that the amino acids of IBF1 (338th Gly, 340th Val, and 374th Arg) important to maintain its function and straw-white hull color (Xia *et al*., 2016; Li *et al*., 2025) are not directly involved in the interaction with CHS1 but affect the structure of the interaction surface with CHS1. These results support our model and suggest that the impairment of IBF1-CHS1 interaction is a common mechanism leading to dysfunctions of IBF1 and enhanced flavonoid accumulation via increased stability of CHS1.

The allele mining analysis suggested that mutations causing dysfunction of IBF1, similar to DJ123, are exclusively distributed among South Asian countries, largely within the *aus* subpopulation (Table S4). The presence of diverse sequence variations at *IBF1* may suggest that *IBF1* locus may have been selected multiple times during the long history of rice cultivation, similar to what occurred for the gibberellin oxidase gene, *OsGA20ox* (also known as *SD1*) determining plant height (Asano et al., 2011). The exclusive distribution of the 1-bp and 2-bp deletions within South Asia may suggest that these alleles have been selected in South Asia, while other variants, such as the R374C, may have been selected in other locations to provide diversity in hull color. This raises the possibility that hull color was an agronomically important trait for ancient farmers. Supporting this, the nucleotide diversity near the *IBF1* locus is low in domesticated rice compared to the wild rice population (Li *et al*., 2025).

Purple-black hull color observed in some rice accessions carrying the 1-bp or 2-bp nucleotide deletions (Table S4) could be due to a combination of dysfunctional IBF1 and functional *A1* encoding a dihydroflavonol reductase (Sun et al., 2018), which converts dihydroflavonols into leucoanthocyanidins, leading to increased anthocyanin synthesis. Thus, a combination of allelic variation at IBF1 and other loci, such as *A1* or other crucial genes (such as *C1* encoding an R2R3-MYB transcription factor and *S1* encoding a bHLH transcription factor) (Sun et al., 2018), may add to the diversity in hull color and its metabolism in a wide genetic pool of rice.

### Effects of hull-derived flavonoids on bacteria

Certain kinds of flavonoids, including luteolin, disrupt iron homeostasis of bacteria by reducing Fe^3+^ to Fe^2+^ and downregulating iron uptake (Zhong et al., 2023). Thus, it is likely that hull-derived flavonoids interfere with physiological functions of bacteria related to iron uptake and usage. *Pantoea* strains utilized in the current study are pathogenic or beneficial endophytes colonizing the apoplast (Dutkiewicz et al., 2016). Thus, a complex interaction and competition in the apoplast including flavonoid accumulation, changes in iron availability and usage by bacteria and plants may occur. Further investigations are necessary to identify the causal metabolite that suppresses siderophore-producing activity, as well as the mode of action.

Apigenin, one of the metabolites of which content was significantly increased in CSSL #14, exhibits antimicrobial activity by inhibiting DNA gyrase activity in a pathogenic microbial species *Staphylococcus aureus* (Morimoto *et al*., 2023). Thus, flavonoid-enriched hull extracts from CSSL #14 conferred by IBF1_DJ123_ may provide a source for natural antibacterial substances. Supporting this, a GWAS study showed that a marker near *IBF1* (7.17 MB on chromosome 9) is closely associated with genotypic differences in seed-derived antimicrobial capacity in rice (Yoshida *et al*., 2022). On the other hand, flavonoids are an important component of plant root exudates (Cesco *et al*., 2010) and attract beneficial microbes to plant roots. Apigenin released from maize roots is a key factor to establish colonization of beneficial bacterial taxa belonging to the Oxalobacteraceae family, thereby promoting plant growth under nitrogen-limited conditions (Yu *et al*., 2021). Other studies also demonstrated the effectiveness of root-originated apigenin and luteolin in attracting nitrogen-fixing beneficial microbes through metabolic engineering of rice and wheat (Yan et al., 2022; Tajima et al., 2025). The specific expression of *IBF1* in hulls (Huang et al., 2012) (Fig. S13) may offer an ideal source of protection against pathogens in seeds without affecting the composition of root exudates. It would be tempting to investigate if IBF1_DJ123_ provides pathogen resistance in field level and if hull-derived metabolites could improve the composition of root-associated microbes and plant performance.

### Conclusions

In conclusion, we propose the mechanism (Fig. 7) in which differences in the allelic state of IBF1 lead to changes in metabolite accumulation patterns and physiological functions of rice hulls. Further metabolite fractionation analysis would be necessary to gain insights into the causal flavonoids that suppress bacterial growth and siderophore generation activity, as well as investigating the possibility of the application of flavonoid-enriched hull extracts in agriculture.

**Figure 7.**
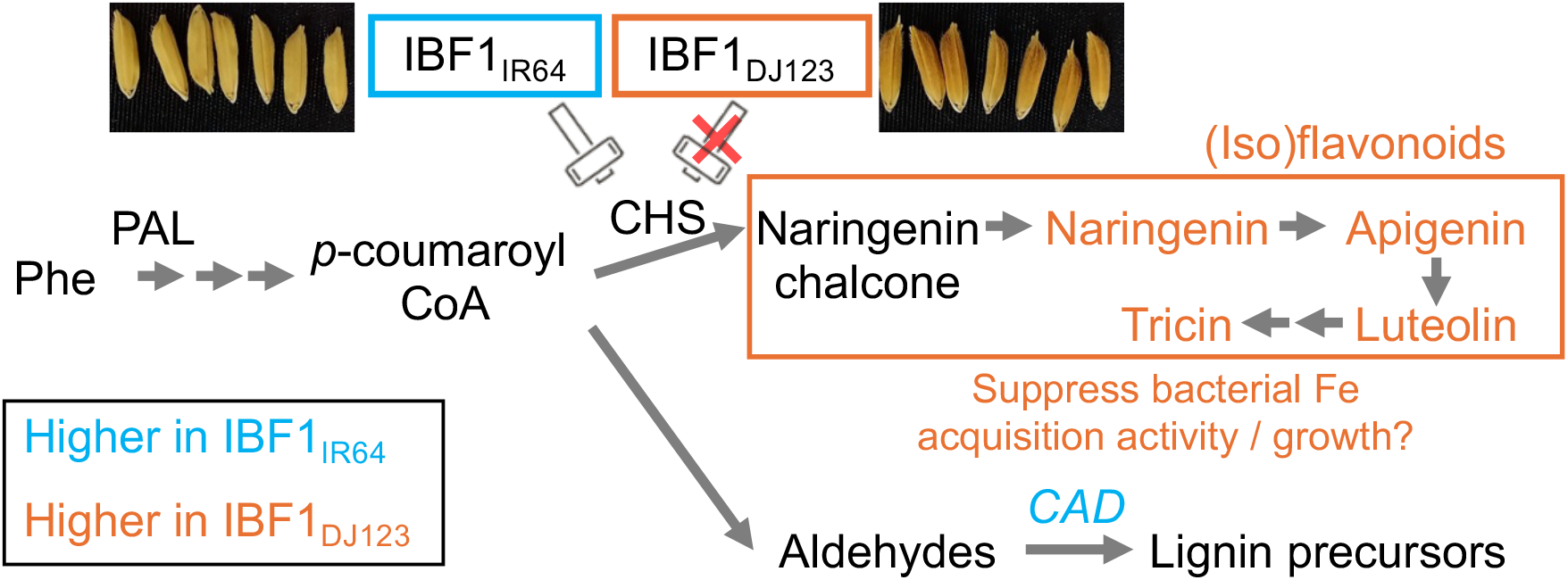
Proposed model of change in metabolism and functionality of rice hulls by different alleles of IBF1. IBF1_IR64_ can interact with and degrade CHS1, while the domain deletion in IBF1_DJ123_ results in the impairment of the IBF1-CHS1 interaction, maintaining the abundance of CHS1. In plants carrying the IBF1_DJ123_ allele, the substrate of CHS1, *p*-coumaroyl CoA, flows into the flavonoid synthesis pathway. The increased supply of the upstream metabolite in the flavonoid synthesis pathway (naringenin chalcone) likely accelerates the synthesis of diverse flavonoids. Conversely, in plants carrying the IBF1_IR64_ allele, *p*-coumaroyl CoA is utilized for the synthesis of aldehydes, potentially leading to an increase in the expression of *CAD*, which is involved in the synthesis of lignin precursors. The increased flavonoid content in plants with the IBF1_DJ123_ allele suppresses bacterial siderophore producing activity and growth. CAD, cinnamyl alcohol dehydrogenase, CHS, chalcone synthase, PAL, phenylalanine ammonia lyase, Phe, phenylalanine.

## Materials and Methods

### Plant materials

Seeds of the BC_1_F_5_ generation of the IR64/Mudgo//IR64 backcross inbred line population were obtained after 4 cycles of selfing, following a backcross of an F_1_ plant of IR64 x Mudgo progeny to the recurrent parent IR64. Seeds obtained from a single plant of BC_1_F_5_ lines were used for the experiments. Phenotypic observations for hull color and sequencing-based genotyping via RAD-Seq were conducted during the field experiment using BC_1_F_6_ plants.

Another mapping population consisted of recombinant inbred lines developed from IR64 x DJ123 as reported previously (Ueda *et al*., 2024). F_5_ plants of the IR64 x DJ123 population were used for phenotypic observations.

A BC_4_F_6_ generation of chromosomal segment substitution line deriving from the IR64 x DJ123 cross (CSSL #14) was developed after repeated cycles of backcrossing and genotypic selections (Fig. S2). Genotyping of intermediate generations was performed via SNP array, RAD-Seq or Kompetitive Allele-Specific PCR (KASP)-based methods.

### Growth conditions and phenotyping of hull color

The first QTL mapping population (IR64 x Mudgo, BC_1_F_6_ plants) was grown in a lowland condition at the experimental field in the Tropical Agriculture Research Front of JIRCAS located in Ishigaki, Japan, during August-November 2021. The second QTL mapping population (IR64 x DJ123, F_5_ plants) was grown in a lowland condition at the experimental field in JIRCAS located in Tsukuba, Japan, during April-September 2019. In both experiments, the fields were fertilized with NPK following local conventional practice (Ueda *et al*., 2024). After harvesting, seeds were air-dried until a constant weight was achieved, scanned by a scanner and manually evaluated for its coloration using a hull color score that ranges from 0 (no pigmentation) to 3 (strong brown pigmentation).

The fine-mapping population (in 2021) and CSSL for hull color evaluation (in 2024) were grown in the same paddy fields of JIRCAS located in Tsukuba. In the fine-mapping experiment, the hull color was visually classified into 2 categories: no or little pigmentation caused by the IR64 or heterozygous alleles; strong pigmentation caused by the DJ123 allele (Fig. S3) in the field.

Hull samples for RNA-seq were obtained from IR64 and DJ123 (in 2021) to identify metabolic genes abundantly expressed in hulls. Panicles at the early grain filling stage were selected, and hulls were harvested by snap-freezing with liquid nitrogen. Hull samples for RNA-seq were obtained from IR64 and CSSL #14 (in 2024) 17 d after heading (DAH) to identify differently expressed genes. The hulls of IR64 and CSSL #14 were also harvested at 0 and 46 DAH for metabolite analyses. The samples were stored at -80°C until the analysis.

Hull samples for total flavonoid and microbial activity analyses were air-dried until a constant weight is reached and stored at room temperature.

### DNA extraction and RAD sequencing

To obtain genome-wide marker data for the IR64 x Mudgo population, DNA was extracted from the leaf of a single BC_1_F_6_ plant (176 individual plants) and 2 replications of parents (IR64 and Mudgo) using a standard phenol-chloroform method, including ethanol precipitation. The library for restriction site-associated DNA sequencing (RAD-Seq) was constructed as reported previously (Kobayashi *et al*., 2017). The library was sequenced by the Hiseq X Instrument (Illumina Inc, CA, USA), generating 150-bp paired-end reads.

### RAD-Seq data analysis and QTL mapping

The analysis of raw sequencing data from RAD-Seq was conducted as reported previously (Ranaivo *et al*., 2022), using the reference genomes of Nipponbare (IRGSP-1.0) (Kawahara *et al*., 2013) and IR64 (GCA_009914875.1) (Zhou *et al*., 2023) (Supplemental Note 1). QTL mapping was conducted using R/qtl essentially as in a previous study (Ueda *et al*., 2024) with the composite interval mapping method using the cim function specifying n.marcovar = 4 option. The significance threshold was determined by 100 times of permutation. QTL mapping using the IR64 x DJ123 population was performed in the same manner as above using the marker data for 654 SNPs (Ueda *et al*., 2024).

### Whole genome sequencing and sequence data analysis

The genomic DNA obtained from Mudgo was subjected to whole-genome re-sequencing using HiSeq X instrument (Illumina), generating 150-bp paired-end reads. Obtained reads were processed essentially as reported previously (Pariasca-Tanaka *et al*., 2025), using the Nipponbare (IRGSP-1.0) (Kawahara *et al*., 2013) reference genome. The genome sequence read for IR64 was obtained from the NCBI SRA database (SRX26568225) (Dinh *et al*., 2025). Extracted variants were visualized in Tassel5 software (Bradbury *et al*., 2007) and converted to hapmap format.

### RNA extraction and RNA sequencing

Hulls were isolated from frozen seeds using tweezers on dry ice. RNA was extracted from pulverized frozen hulls using the ISOSPIN Plant RNA kit (Nippon Genetics) following the manufacturer’s instructions, including DNase treatment. The library for RNA sequencing was constructed using the NEBNext Ultra II Directional RNA Library Prep Kit for Illumina (New England Biolabs, Ipswich, USA) (n=2 for IR64 and DJ123 hulls comparison) or VAHTS Universal V8 RNA-seq Library Prep Kit for Illumina (Vazyme, Nanjing, China) (n=3 IR64 and CSSL #14 hulls comparison) according to the manufacturer’s instructions. The library was sequenced with the NovaSeq 6000 Instrument (Illumina, San Diego, USA), generating 150-bp paired-end reads. The raw data analysis followed a previous report (Ueda, 2024) (Supplemental Note 1).

Differentially expressed genes were determined by the R software DESeq2 (Love *et al*., 2014) using count data with the nbinomLRT command. Genes with adjusted *P* values of < 0.05 were considered significant.

### Yeast two-hybrid assay

The yeast two-hybrid assay using *Saccharomyces cerevisiae* strain Y2H Gold (Takara Bio Inc., Kusatsu, Japan) was conducted as previously reported (Ishizaki *et al*., 2023), using pGBKT7 (for prey proteins) and pGADT7 (for bait proteins) vectors carrying inserts amplified with gene-specific primers containing restriction sites (Table S5). Transformed cells were initially selected on synthetic defined (SD) medium plates lacking leucine and tryptophan (SD/-LW). A single colony was streaked onto SD/-LW medium or SD medium plates lacking leucine, tryptophan, histidine, and adenine (SD/-LWHA). The plates were incubated at 30°C and photographed after 5 d.

### Metabolite analyses

Total flavonoid content was measured according to a previous report (Shao *et al*., 2012) using air-dried hull samples. Hulls isolated from seeds were ground into a fine powder with liquid nitrogen and mixed with 10-fold volume of 80% methanol. After incubating at 37°C for 30 min, the supernatant was obtained after 5 min of centrifugation at 12,000 x g, mixed with a 1/9 volume of 10% AlCl_3_, and incubated for 30 min at room temperature. The absorbance was then recorded at 420 nm. The reaction containing 80% methanol instead of AlCl_3_ solution was used as the baseline for each sample.

Metabolome analysis was conducted using extracts obtained with 100% methanol and liquid chromatography-mass spectrometry (LC-MS) (Supplemental Note 1). The obtained raw data were analyzed using the PowerGetBatch software as reported previously (Sakurai et al., 2023). MS/MS spectra were obtained from 744 metabolite peaks in positive ionization mode. A total of 46 potential flavonoid peaks were annotated through the analysis of MS/MS spectra, as determined by the FlavonoidSearch system (Akimoto *et al*., 2017), with a score of ≥ 0.05. The structure of the compound was further predicted by the MASST (Wang et al., 2020a) and MS-FINDER (Tsugawa et al., 2016). The metabolite peaks for luteolin, apigenin/genistein, Tyr, Phe, Ile, and Leu were identified at the annotation level 1 (Blaženović et al., 2018) by comparing the *m/z* values, retention times, and MS/MS spectra with those obtained from authentic standard compounds using the same LC-MS conditions.

### Protein modeling via Alphafold3

Amino acid sequences of different versions of IBF1 and CHS1 were subjected to an online server of Alphafold3 (Abramson *et al*., 2024) (https://alphafoldserver.com) with default settings. The output files of Alphafold3 were further analyzed by the Predictome website (https://predictomes.org/tools/af3/) (Schmid and Walter, 2025) with default settings to identify pairs of amino acids that potentially mediate the interaction between two proteins.

### Growth and physiological activity assays in bacteria

Metabolites in air-dried hulls were extracted using 80% methanol at a rate of 40 mL/gDW at 37°C for 30 min. The supernatant was collected after centrifugation at 12,000 x g for 10 min at 4 °C. A 5-mL aliquot was freeze-dried, and the pellet was resuspended in 2.0 mL of sterile deionized water.

Bacterial species (*Pantoea agglomerans* 102470, *Pantoea* strain 6PN2, *Staphylococcus pasteuri* 105, *Staphylococcus warneri* 108) were pre-cultured in 3 mL of Trypticase soy broth (TSB) at 25 °C for 16-20 h. Following centrifugation, the bacterial pellets were washed three times with 10 mM MgCl_2_ and resuspended in the same buffer to achieve OD_600_ = 0.025. The bacterial suspension was mixed with an equal volume of hull extracts or deionized water and incubated at room temperature for 16-18 h. Activities for cellulase, phosphorus solubilization, and siderophore production were evaluated by dropping the bacterial resuspension on agar plates containing sodium carboxymethyl cellulose (Liang et al., 2014), phosphorus medium (Chen and Liu, 2019), or chrome azurol S (Louden et al., 2011) and incubating in dark at room temperature. Growth assay was conducted by inoculating 150 µL of a 100-fold diluted bacteria-hull extract mixture (in TSB medium) in a 96-well microplate and monitoring OD_600_ at 25 °C for 48 h.

### Database analysis

The number of genotypes and allele frequency of *IBF1* variant was analyzed on European Variant Archive (EVA) database (https://www.ebi.ac.uk/eva/; accessed on 25th August, 2025). The hull color of the selected accessions was visually evaluated using photos retrieved from the Genesys database (https://www.genesys-pgr.org/; accessed on 26th August, 2025).

### Statistical analyses

The results of targeted metabolite analysis were analyzed via two-way ANOVA using VassarStats online calculator (http://vassarstats.net/anova2u.html) to evaluate the effect of genotype and developmental stage. Comparison of flavonoid content (Fig. 1E) and effects of hull extracts on physiological activities of microbes were compared by Student’s *t*-test.

## Accession Numbers

The raw data for RNA-seq, RAD-seq and whole-genome sequencing have been deposited in NCBI SRA database under the accession numbers PRJNA1318644 (RNA-seq for the comparison of IR64 vs DJ123 hulls), PRJNA1321561 (RNA-seq for the comparison of IR64 vs CSSL #14 hulls), PRJNA1322785 (RAD-seq for the IR64 x Mudgo population), and PRJNA1320712 (whole-genome resequencing of Mudgo). The metabolome data have been deposited in Metabolights database under the accession number MTBLS12978.

## Supplemental Data

**Supplemental Figure S1** | Distribution of 333 SNP markers for the BC_1_F_6_ population derived from IR64 and Mudgo.

**Supplemental Figure S2** | Development history of CSSL #14.

**Supplemental Figure S3** | Appearance of seeds with different IBF1 alleles.

**Supplemental Figure S4** | Sequence variation of IBF1.

**Supplemental Figure S5** | Results of marker analysis within the anticipated deletion region in Mudgo.

**Supplemental Figure S6** | Negative controls for the yeast two-hybrid assays.

**Supplemental Figure S7** | Neither IBF1_IR64_ nor IBF1_DJ123_ interacts with phenylalanine ammonia lyases.

**Supplemental Figure S8** | 3D structure modeling of IBF1_DJ123_-CHS1_IR64_ interaction.

**Supplemental Figure S9** | Potential effects of other IBF1 mutations on the formation of IBF1-CHS1 complex.

**Supplemental Figure S10** | Coloration of reaction mixture for flavonoid analysis.

**Supplemental Figure S11** | Time-course analysis of selected amino acids in hull extracts.

**Supplemental Figure S12** | Physiological and growth analyses using hull extracts in *Staphylocossus* species.

**Supplemental Figure S13** | Expression of *IBF1* in various parts of Nipponbare plants.

**Supplemental Table S1** | Detected peaks in untargeted metabolome analysis.

**Supplemental Table S2** | Candidate peaks of flavonoid aglycones detected differently in IR64 and CSSL #14.

**Supplemental Table S3** | Differentially expressed genes in hulls of IR64 and CSSL #14.

**Supplemental Table S4** | Distribution of 1-bp and 2-bp nucleotide deletions similar to DJ123.

**Supplemental Table S5** | Primers used in this study.

**Supplemental Note 1** | Additional Methods.

## Acknowledgments

The authors thank Dr. A.Z. Htwe, M. Yonemoto, K. Nishihara and M. Matsuyama for assistance in DNA extraction and plant phenotyping. The authors also thank field technicians of JIRCAS for management of plants in the field.

## Author Contributions

YU conceptualized the study, performed plant phenotyping, DNA extraction, genotyping, QTL mapping, fine-mapping, RNA sequencing, protein-protein interaction assays, protein structural analyses, total flavonoid assays and sequence analyses, interpreted the data and prepared manuscript draft and figures and tables. YU, TI designed the experiments. YU, JP-T, KK and MW developed genetic materials. JP-T and MW performed whole-genome resequencing of Mudgo. YM performed bacterial assays. NS performed metabolome analyses and analyzed these data. YU, HS, JP-T, KK and MW conducted field experiments. HT constructed RAD-seq library. NS and MW provided editorial comments. All authors have checked the final version of the manuscript and agreed to submission.

