## Supplementary material for "Natural variation in IBF1 disrupts its interaction with CHS1 and affects metabolism of hulls in rice": Fig. S1

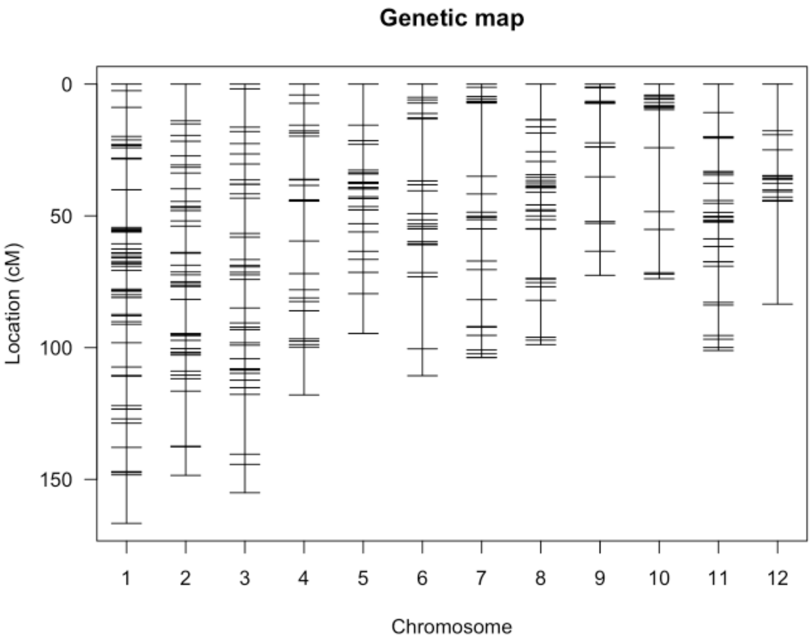


**Fig. S1 | Distribution of 333 SNP markers for the $BC_1F_6$ population derived from IR64 and Mudgo.**

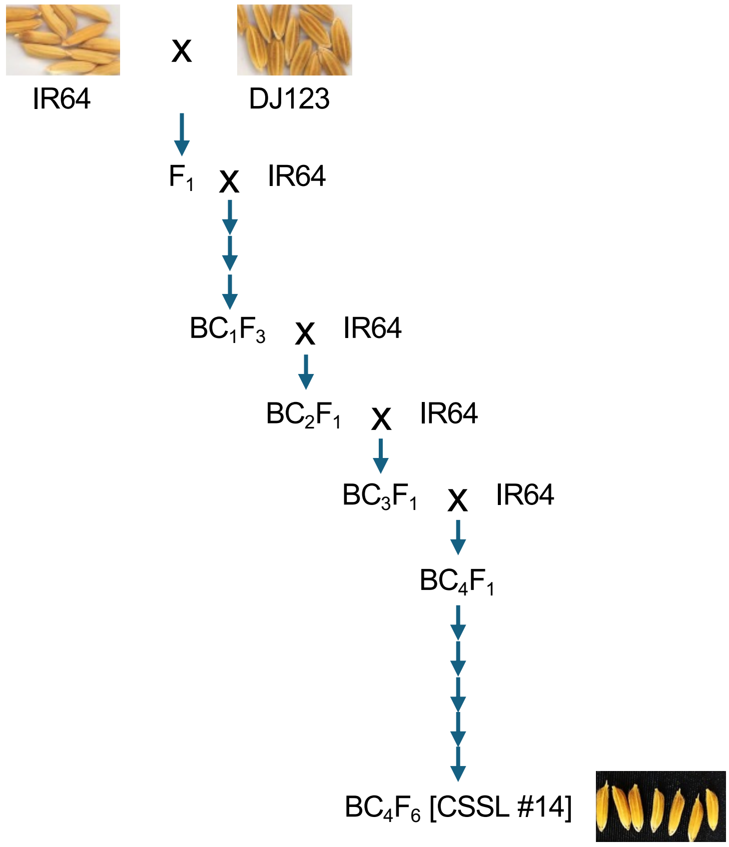


**Fig. S2 | Development history of CSSL #14.**

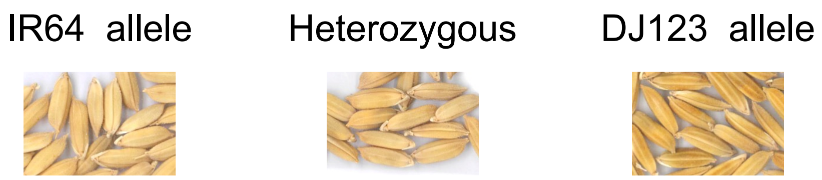


**Fig. S3 | Appearance of seeds with different IBF1 alleles.**

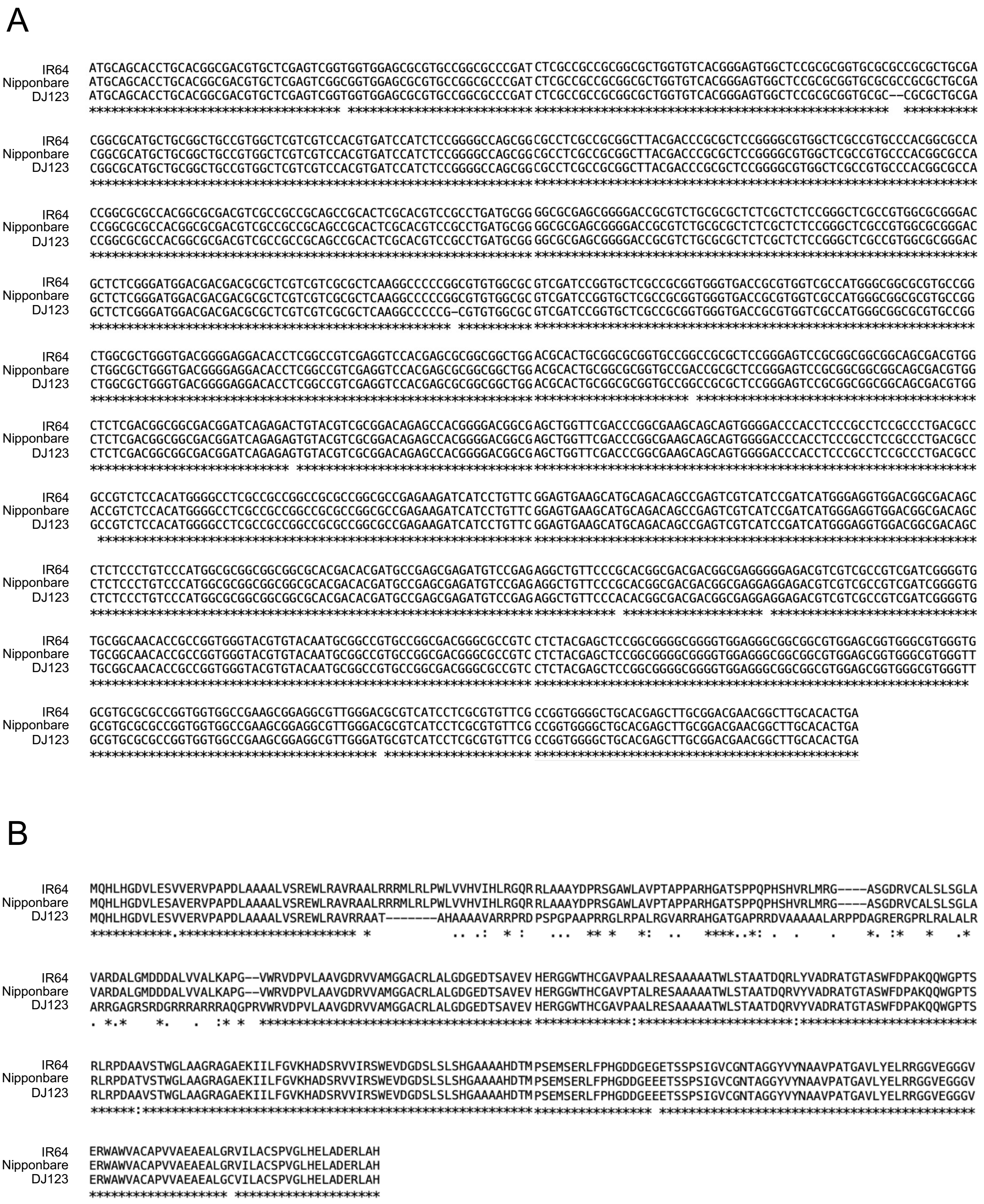


**Fig. S4 | Sequence variation of IBF1.**

The results of the alignment of the nucleotide sequence of *IBF1* (A) and the amino acid sequence of IBF1 (B) in Nipponbare, IR64, and DJ123 are shown.

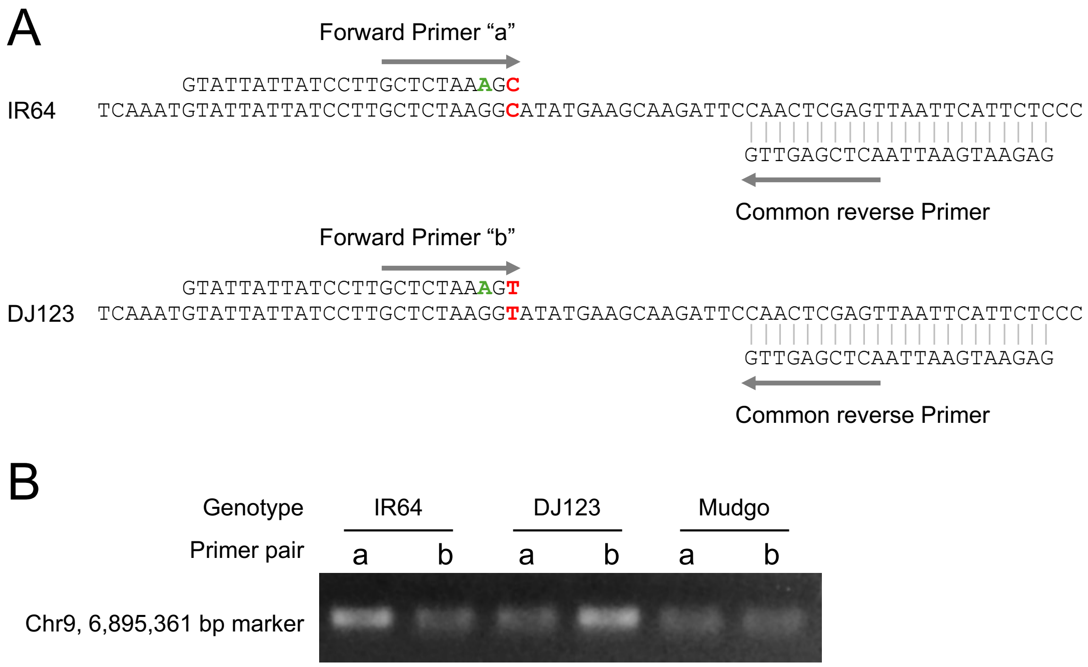


**Fig. S5 | Results of marker analysis within the anticipated deletion region in Mudgo.**

(A) Details of allele-specific marker at 6,895,361 bp on chromosome 9. The red and green bases represent the target SNP marker and a mismatch base added to the primer to minimize non-specific amplification (Liu et al. 2012, *Plant Methods* 8, 34; Pariasca-Tanaka et al. 2025, *Curr Plant Biol* 42, 100469). (B) Results of the marker analysis using the primers shown in A.

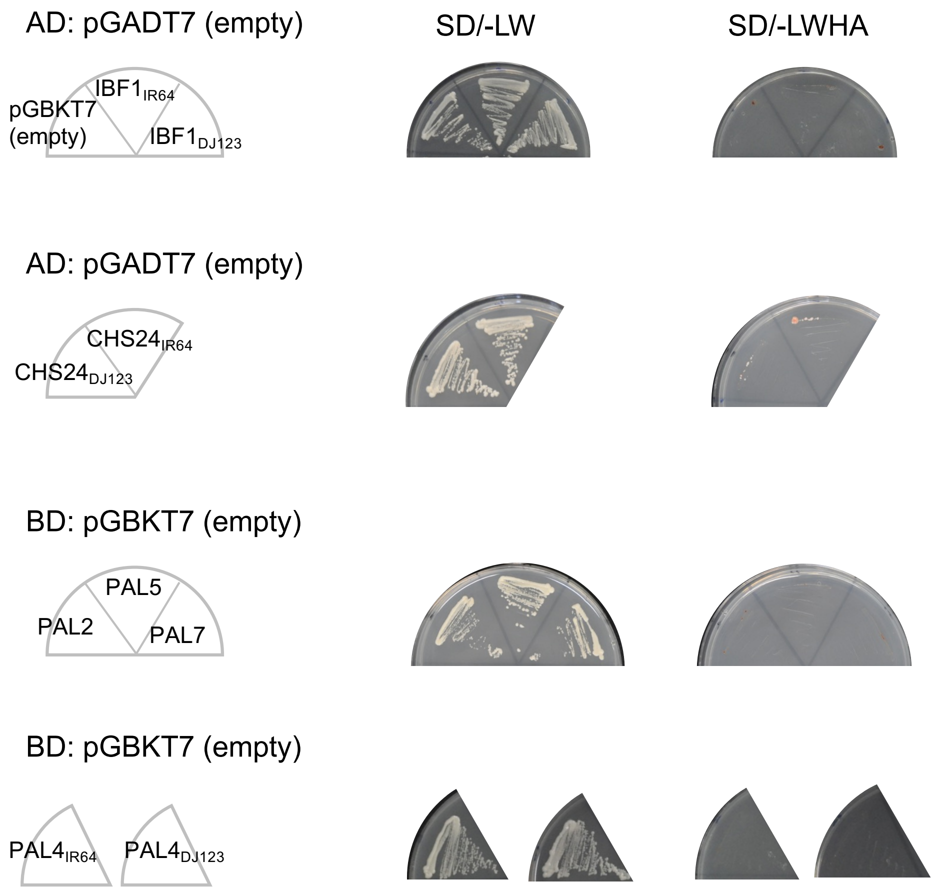


**Fig. S6 | Negative controls for the yeast two-hybrid assays.**

SD/-LW and SD/-LWHA repsresent synthetic defined (SD) media lacking Leu and Trp (-LW) or Leu, Trp, His and Ade (-LWHA).

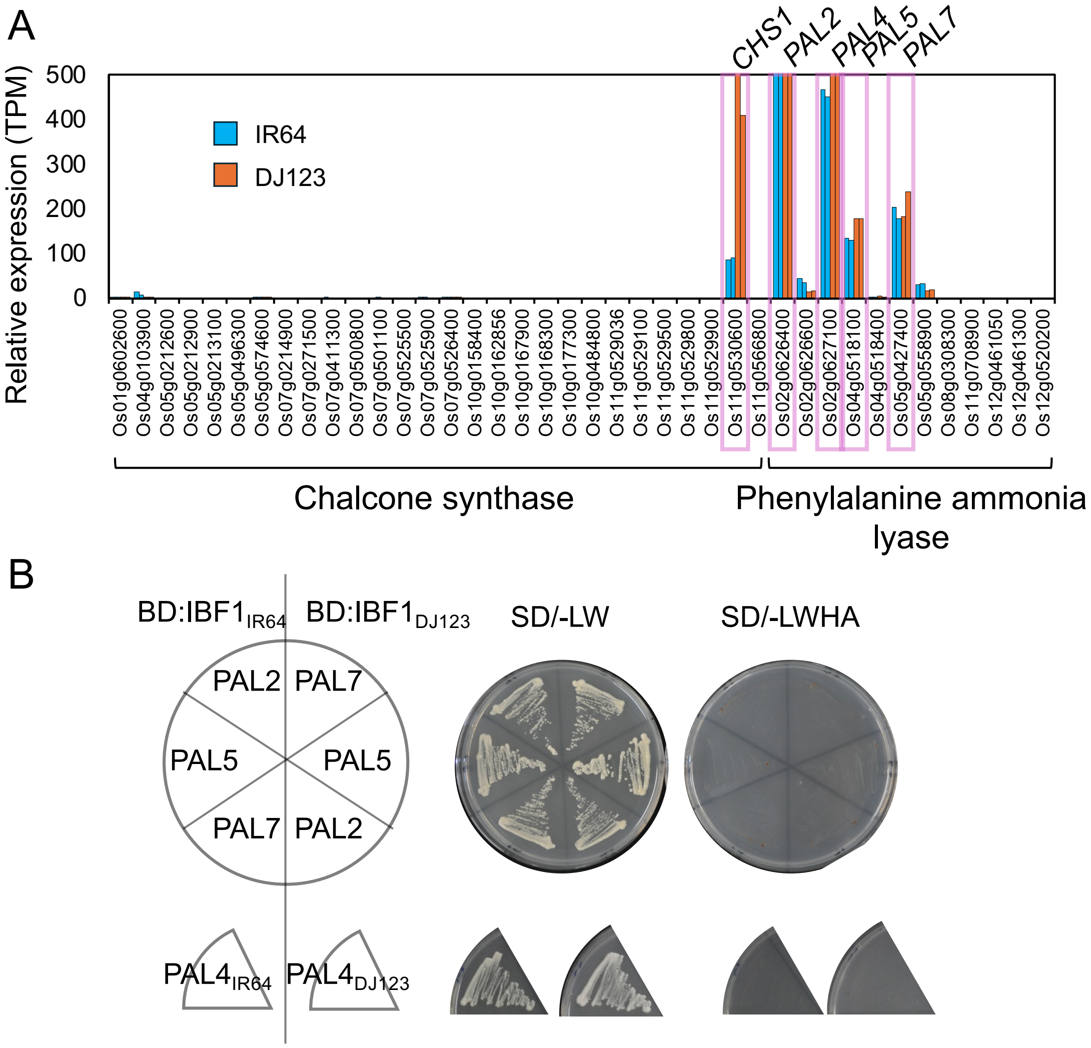


**Fig. S7 | Neither IBF1$_{IR64}$ nor IBF1$_{DJ123}$ interacts with phenylalanine ammonia lyases.**

(A) Expression of all chalcone synthases and phenylalanine ammonia lyase genes in young hulls of IR64 and DJ123.

(B) Results of yeast two-hybrid assays. The left panel shows the combination of bait (IBF1) and prey (PAL) proteins used for the assay. The amino acid sequence for PAL4 was different for IR64 and DJ123, and both were examined. The right panel shows the growth of yeasts introduced with the respective combinations of proteins. SD/-LW and SD/-LWHA indicate selection synthetic defined (SD) media lacking Leu and Trp (-LW) or Leu, Trp, His, and Ade (-LWHA).

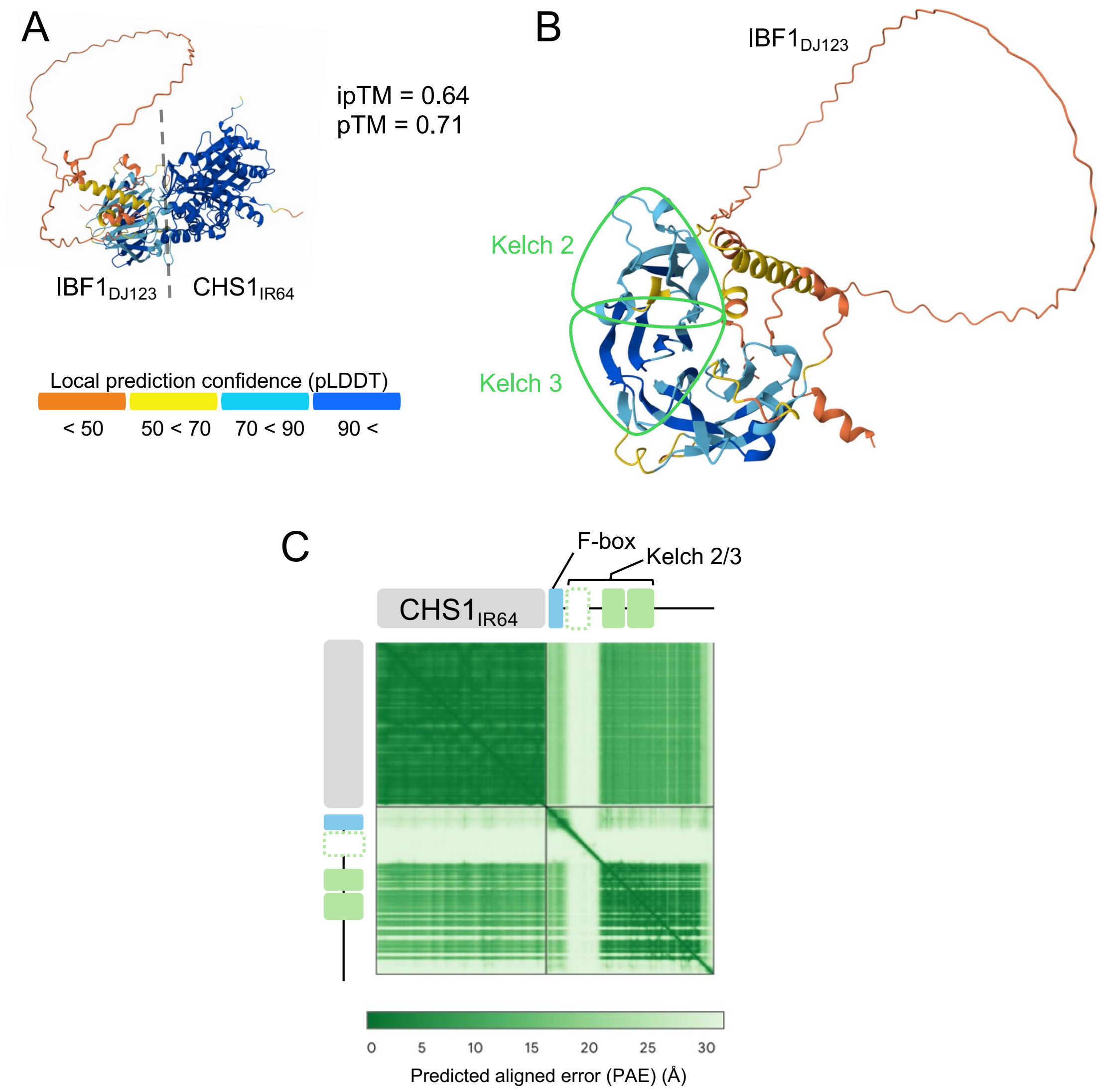


**Fig. S8 | 3D structure modeling of IBF1$_{DJ123}$-CHS1$_{IR64}$ interaction**

(A) Structural modeling of IBF1$_{DJ123}$-CHS1$_{IR64}$. (B) The interaction surface of IBF1$_{DJ123}$ as viewed from the CHS1 side. The CHS1 molecule is omitted from visualization for simplicity. In A and B, amino acid residues are colored based on the accuracy level of prediction judged by pIDDT values. (C) Expected position error for the modeling results of IBF1$_{DJ123}$-CHS1$_{IR64}$.

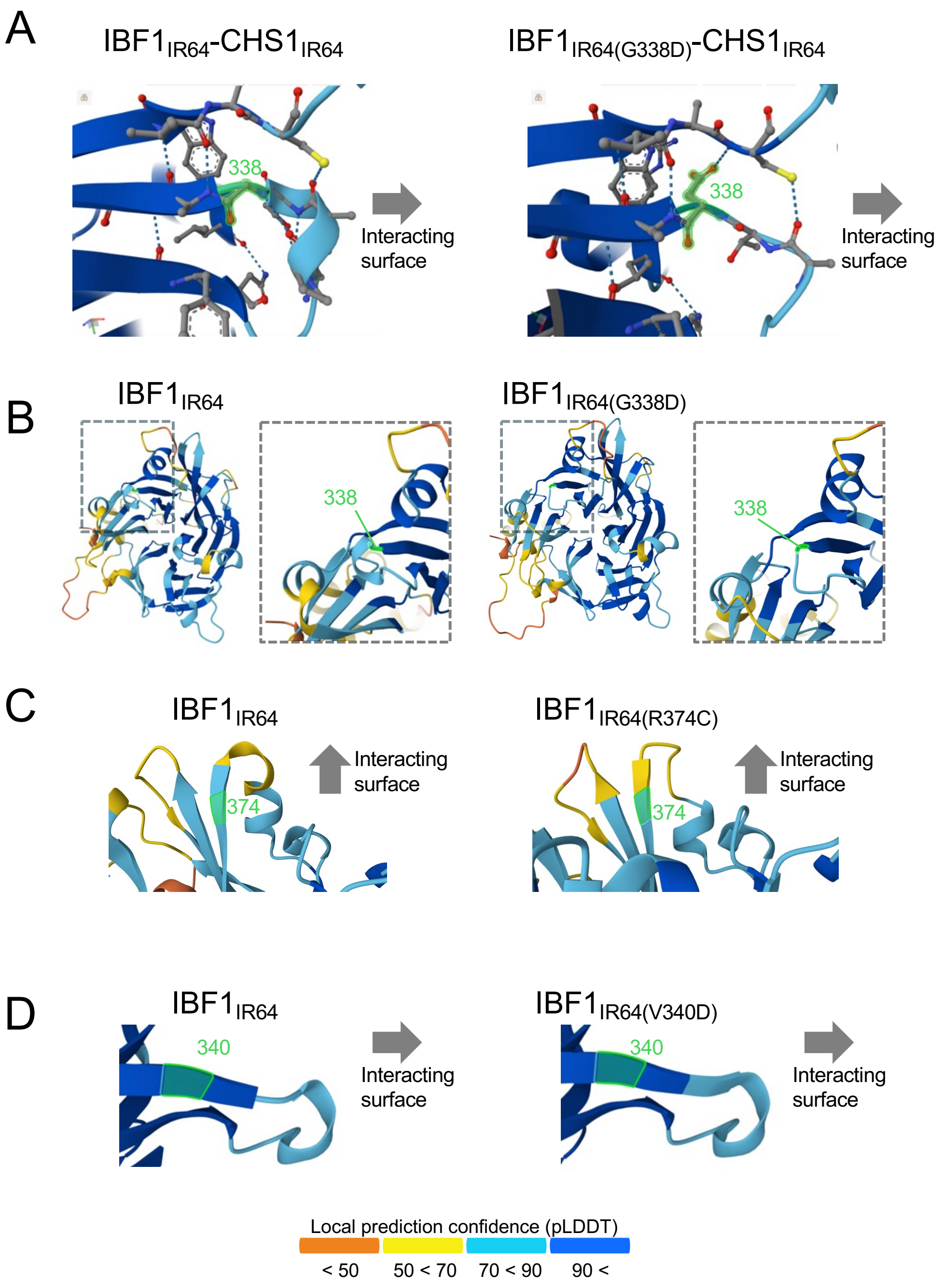


**Fig. S9 | Potential effects of other IBF1 mutations on the formation of IBF1-CHS1 complex.** Structural modeling of IBF1$_{IR64}$-CHS1$_{IR64}$ in wild-type IBF1 and its comparison with G338D (A,B), R374C (C), and V340D (D) versions of IBF1 mutants. In all cases, structural modeling was conducted including CHS1, but CHS1 is omitted from the visualization for simplicity. In A, C, and D, the grey arrows indicate the direction in which CHS1 is positioned. In B, the interaction surface of IBF1 is visualized from the CHS1 side.

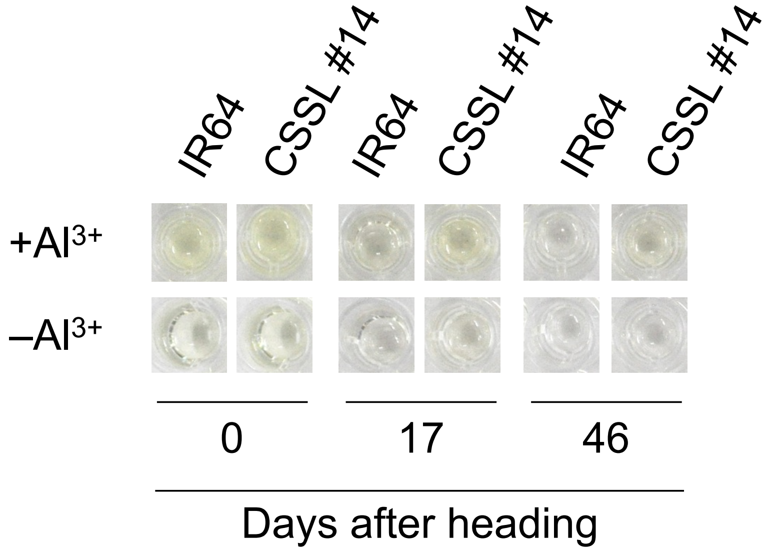


**Fig. S10 | Coloration of reaction mixture for flavonoid analysis.**

The colors of hull extracts using 80% methanol, with and without a final concentration of 1% of $AlCl_3$, are shown.

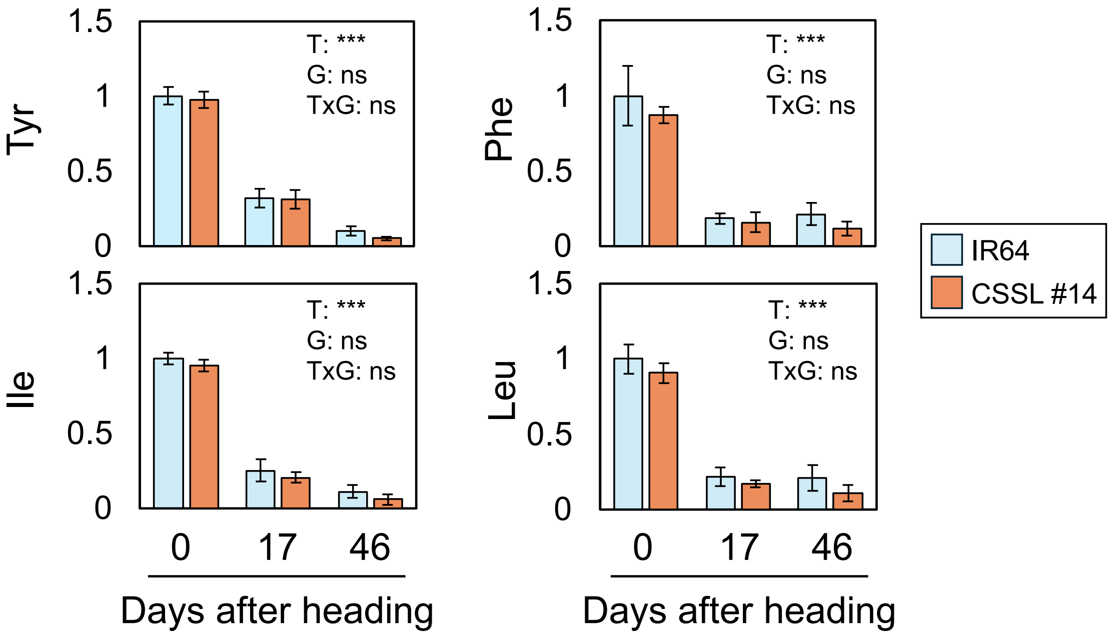


**Fig. S11 | Time-course analysis of selected amino acids in hull extracts.**

Values are means ± S.D. (n=3). Two-way ANOVA was conducted for each amino acid to analyze the effect of time (T), genotype (G) and their interaction (TxG), and the results are shown as asterisks (***) for $P < 0.001$. "ns" indicates $P > 0.05$.

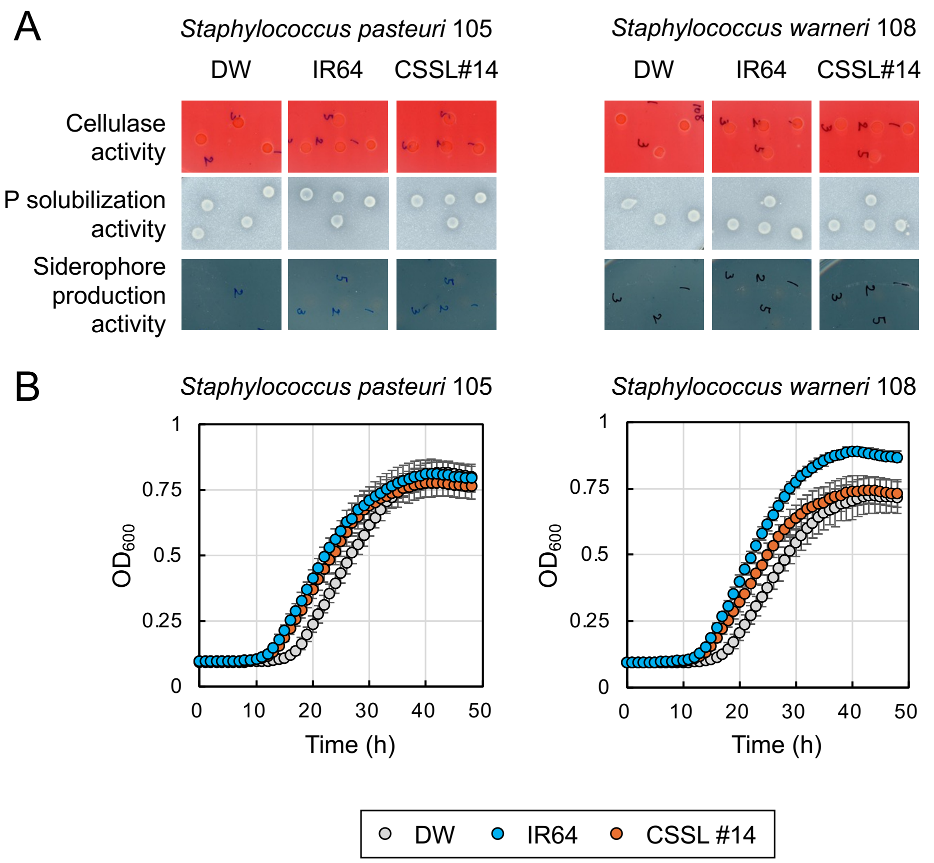


**Fig. S12 | Physiological and growth analyses using hull extracts in *Staphylococcus* species.**

(A) Effects of hull extracts from IR64 and CSSL #14 on physiological characteristics of *Staphylococcus pasteuri* strain 105 and *Staphylococcus warneri* strain 108. (B) Growth analysis of respective strains in the presence of hull extracts from IR64 and CSSL #14. In B and C, values are means ± S.D. (n=8).

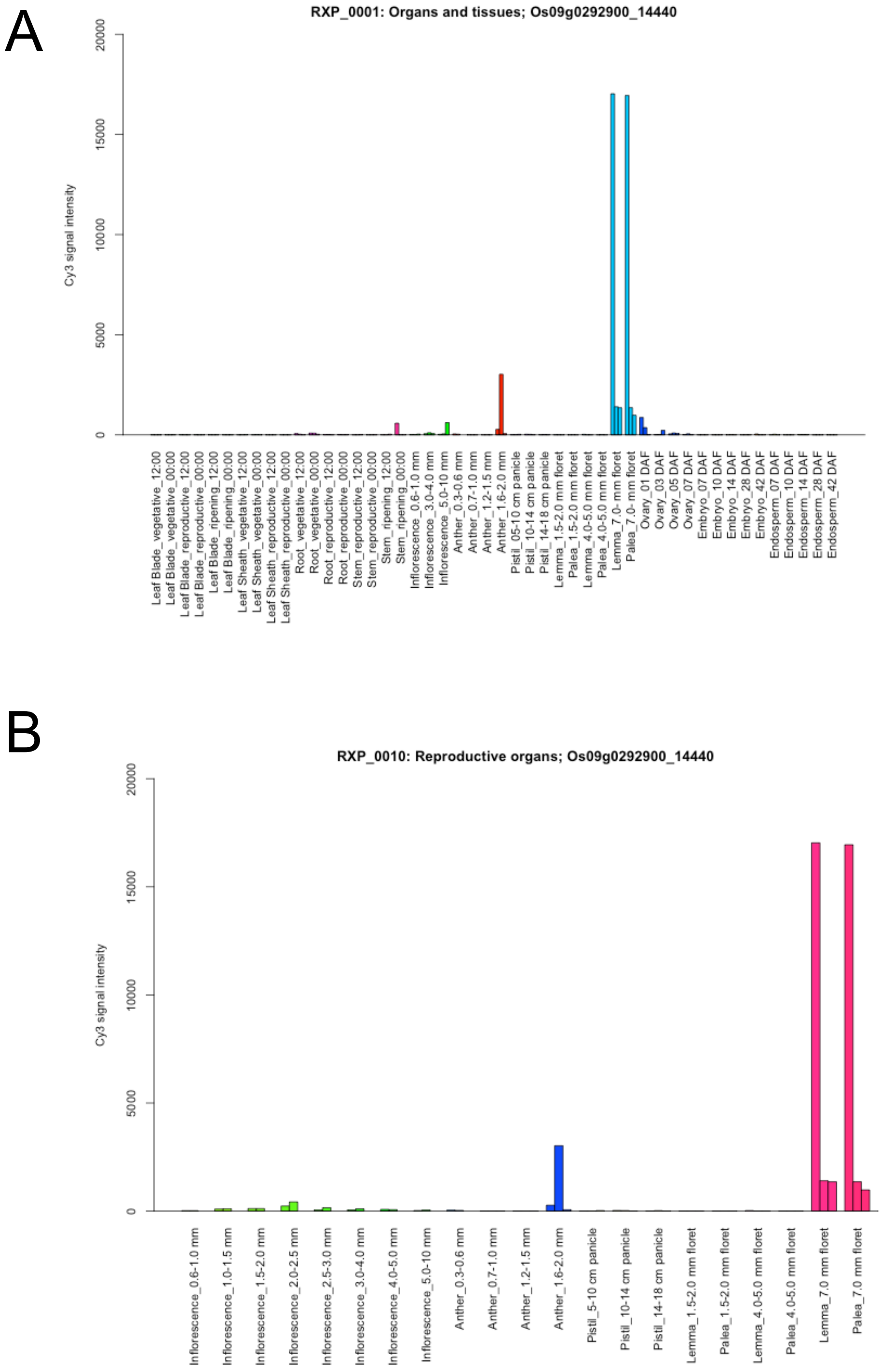


**Fig. S13 | Expression of *IBF1* in various parts of Nipponbare plants.**

(A,B) The expression of *IBF1* in different parts of the whole plant (A) and reproductive organs (B) is shown. Data was obtained from the RiceXPro database (https://ricexpro.dna.affrc.go.jp/) (Sato et al. 2011, *Nucleic Acids Res* 39, D1141–D1148; Sato et al. 2011, *BMC Plant Biol* 11, 10).

**Supplemental Note 1 | Additional Methods.**

*RAD-Seq data analysis*

The quality of raw reads was checked using FastQC software, followed by adapter removal and read trimming using trimmomatic (Bolger et al., 2014) with the following settings: LEADING:3 TRAILING:3 SLIDINGWINDOW:4:15 MINLEN:100. The trimmed reads were then mapped to reference genomes using bwa (Li and Durbin, 2009) and converted to bam format using samtools (Li et al., 2009). After removing reads with low quality (< 60), variants were extracted using bcftools (Danecek et al., 2021) to generate a vcf formatted file. Variants were filtered using vcftools (Danecek et al., 2021) with the following settings: -max-missing 0.95 --min-meanDP 10 --max-meanDP 100 --minQ 20 --min-alleles 2 --max-alleles 2.

SNPs within 100 bp from detected INDELs were further removed using bcftools. To minimize spurious heterozygous calls, a heterozygous allele was defined only when the genotype call was supported by at least 6 reads from each parental allele. The genotype call was otherwise converted to missing. The genotype table was then imputed using LinkImpute (Money et al., 2015) in Tassel 5 software (Bradbury et al., 2007). Furthermore, any missing, heterozygous or inconsistent sites within 2 replications of each IR64 and Mudgo were removed. Initially, the reads were mapped to the reference genomes of Nipponbare (IRGSP-1.0) (Kawahara et al., 2013) and IR64 (GCA_009914875.1) (Zhou et al., 2023). The resulting number of markers was 545 and 546 for Nipponbare and IR64 alignments, respectively. Based on the Nipponbare-aligned marker data, large gaps (> 4 MB) were filled with IR64-aligned marker data where possible. After removing 5 samples with > 80 crossovers, marker orders were corrected using the ripple function with error.prob = 0.005 and window = 4 options in the R package R/qtl. Markers within < 0.5 cM distance were further removed, resulting in a total of 333 markers (318 and 15 markers derived from Nipponbare and IR64 alignment, respectively).

*RNA-Seq data analysis*

The quality of the raw reads was checked using FastQC software. Removal of adapter sequences and read trimming were conducted with trimmomatic. Trimmed reads were mapped to the Nipponbare reference transcript (IRGSP-1.0) using HISAT2 software (Kim et al., 2015). Transcript per million (TPM) values and count data were obtained using StringTie software (Pertea et al., 2015).

*Metabolome analysis*

Metabolome analysis was conducted according to the previous report (Sakurai et al., 2023) with slight modifications as briefly described below. 3-fold volumes of 100% ethanol to the weight of frozen pulverized samples were added, and the mixture was shaken twice with a mixer mill (MM 400, Verder Scientific, Vleuten, Netherlands), each for 2 min at 1500 rpm. The supernatant was obtained after centrifugation at 17,400 x g for 5 min at 4°C. The resulting extract was passed through a C18 solid-phase column (MonoSpin C18, GL Science, Tokyo, Japan) to remove hydrophobic compounds and filtered with a 0.2 μM polytetrafluoroethylene membrane (Millipore, Burlington, USA). Compounds in the extract were separated by the Nexera X2 (Shimadzu, Kyoto, Japan) ultra-high performance liquid chromatography equipped with the InertSustain AQ-C18 column (GL Science) and analyzed with the Compact (Bruker, Billerica, USA) quadrupole time-of-flight mass spectrometry. The column temperature was maintained at 40°C, and the eluent was analyzed with a gradient of 0.1% (v/v) formic acid and 100% acetonitrile at a flow rate of 0.2 mL/min. The release condition for Active Exclusion for MS/MS analysis was set to 0.2 min. The valid peaks were defined as those detected in the rice samples and not or weakly detected in the mock control samples with less than 1/10 intensity of the rice sample, estimated by $\log_{10}$-transformed and median-centered values. Data quality was ensured by the consistency of signals of an internal standard, 1 μM 7-hydroxy-5-methylflavone.

The candidate 46 flavonoid aglycones (Supplementary Table S2) may contain those artificially generated by in-source fragmentation from their derivatives, such as glycosides. Therefore, we examined this by comparing the *m/z* value of candidate aglycones with those of the MS/MS product ion peaks obtained from the other peaks co-eluted and detected within 0.2 min retention time (RT) tolerance. Among 46 candidates, we found only one apparent case of this. The peak number 2702 (RT 14.03 min, *m/z* 301.0679) was co-eluted with a candidate precursor, peak number 2701 (RT 14.03 min, *m/z* 463.1204) which had MS/MS spectrum composed almost entirely of a single product ion with *m/z* 301.0679. Based on the neutral loss mass of the product ion (162.0515), the peak 2701 was annotated as a potential glycoside.
